# Younger is Better But Only for Males: Social Behavioral Development Following Juvenile Traumatic Brain Injury to the Prefrontal Cortex

**DOI:** 10.1101/2025.05.24.655898

**Authors:** Sophia Shonka, Michael J. Hylin

## Abstract

Juvenile traumatic brain injury (jTBI) is associated with persistent social impairments, particularly when injury occurs early in development. The prefrontal cortex (PFC) is especially vulnerable to injury due to its late maturation and location, and developmental disruptions during key phases such as synaptic pruning or myelination may result in long-term behavioral deficits. This study investigated how the age at injury and biological sex influenced the development of social behavior and frontal cortical plasticity. Using a controlled cortical impact model of a bilateral medial PFC (mPFC) injury, we compared injuries sustained on postnatal day (PND) 17 or 28—approximating toddlerhood and middle childhood, respectively. Social behaviors were assessed longitudinally during pre-puberty, puberty, and young adulthood. Play behavior, sociability, social memory, social dominance, and aggression were evaluated, and morphological analyses examined dendritic complexity in the orbitofrontal cortex (OFC) and mPFC using Golgi-Cox staining, and myelin integrity across the mPFC-OFC-amygdala circuit using Luxol-fast blue staining.

We hypothesized that PND 17 injuries would result in greater social deficits and increased aggression compared to PND 28 injuries and shams, with males showing more severe impairments than females. Results partially supported these predictions. While jTBI had no effect on general play engagement, it did alter play initiation: PND 28 injury increased play initiation in both sexes, while PND 17 TBI injury delayed normal play development in females. Injuries had no significant impact on sociability or social memory. However, PND 28 injury increased social dominance and aggression in adulthood, with sex moderating these effects. Specifically, PND 17 injury decreased aggression in males and increased it in females. Childhood play behaviors predicted adult aggression, particularly in PND 28-injured animals, and these relationships were moderated by injury and sex. Despite the behavioral findings, histological analyses revealed no significant group differences in dendritic complexity or myelination, though effect sizes suggested decreased dendritic arborization and increased myelin in PND 28-injured animals.

These findings highlight age- and sex-dependent vulnerability in the development of social behavior following jTBI. Contrary to expectations, early injury had more pronounced effects in females, while later injury was more detrimental for males. The divergence in behavioral outcomes despite limited histological differences suggests complex, possibly circuit-specific, mechanisms underlying these effects. This study underscores the importance of considering both sex and developmental timing in jTBI research and supports the need for longitudinal models to capture evolving behavioral trajectories.

## Introduction

Traumatic brain injury (TBI) is a leading cause of death and disability worldwide (1,2), with particularly severe and long-lasting consequences in children. In the United States alone, over 837,000 injuries are reported annually among children aged 0-14 years (1,3). The consequences of juvenile TBI (jTBI) are often protracted (4), with deficits that evolve or intensify over time as children ‘grow into’ their lesions (5–9). Young children are especially vulnerable to poor outcomes, given that brain injury occurs in the context of dynamic and ongoing development of neural systems (3,10). In particular, damage during critical periods of synaptogenesis (8,9,11), myelination (12), or synaptic pruning (7,9,11,13–16) may derail the trajectory of cortical maturation, compounding functional impairments across time (11,17).

The frontal lobe is of particular concern in pediatric TBI due to its prolonged developmental trajectory (18). Unlike other cortical regions that mature relatively early, the prefrontal cortex (PFC) continues to develop into early adulthood (3,17,19), undergoing extensive synaptic pruning (17,19), myelination (20–22), and connectivity reorganization (23,24). These developmental processes confer both opportunity for recovery (9,25) and heightened vulnerability (8,9,25,26). Disruptions during these windows may have lasting consequences for cognition, emotional regulation, and social functioning (5,27).

Indeed, frontal lobe injury during development is frequently associated with executive dysfunction and social impairments (27–30), which can persist into adulthood (29,30) and interfere with adaptive functioning (3,29). Social deficits, including impaired peer relationships and social cognition, are especially concerning given their role in emotional well-being and long-term mental health (31). Prior work has shown that damage to the medial PFC (mPFC) (32–36), orbitofrontal cortex (OFC) (37,38), and amygdala (37,38)—key regions supporting social behavior (39–41)— can lead to deficits in social memory, empathy, and emotional regulation. However, the developmental timing of such injuries appears to modulate the severity and nature of these impairments (8,42).

In animal models, the effects of TBI during early postnatal development vary depending on the age at injury. For instance, injuries at postnatal day (PND) 17, a time corresponding with peak synaptogenesis (43), lead to significant neuronal loss (42,44,45), altered dendritic complexity (16), and spatial memory deficits (42,46). However, these studies have primarily focused on parietal injuries. While direct investigations of frontal injury at PND 17 are lacking, some inferences can be drawn from studies examining jTBI at PND 21. At this age, frontal lobe injury resulted in no observable deficits in spatial learning, memory, or sociability (47), despite notable cell death at the injury site (47,48) and adjacent tissue (47), as well as evidence of exaggerated synaptic pruning (7).

In contrast, injuries occurring between PND 28 and 30 have been associated with social play deficits and social rejection (49), spatial learning impairments (26,50), cortical thinning (25), and abnormal synaptic pruning (15), although synaptic organization appeared unaffected (51). Despite limited research on frontal injuries specifically at PND 28, the ongoing development of PFC myelination during this window, and the vulnerability of unmyelinated axons to injury (52,53), suggests that damage at this stage may produce substantial behavioral consequences. Overall, while both PND 17 and PND 28 represent critical windows for brain development, it remains unclear which period is more vulnerable to frontal injury.

Moreover, relatively few studies have directly compared the impact of injury timing on the development of social behavior across key developmental milestones (e.g., pre-puberty, puberty, adulthood) (6,12,48). Most preclinical studies have relied on single time-point assessments, limiting insight into how jTBI affects the unfolding of social capacities over time. Even fewer studies have examined how sex interacts with age at injury to shape outcomes (7,49), despite calls from the Institute of Medicine to prioritize sex differences in neuroscience research (54). Evidence from both clinical and preclinical studies suggests that females may be relatively protected from the deleterious effects of TBI (7,55,56), potentially due to hormonal influences (57–60) and structural differences in white matter organization (61,62). Yet, direct comparisons across sex and injury age in a single design remain scares.

Given this gap, there is a critical need for longitudinal, developmentally informed animal models that investigate how age at injury and biological sex jointly influence social behavioral trajectories and associated neural substrates. In particular, frontal circuits—including the mPFC and OFC—warrant focused examination given their centrality to social functioning and known sensitivities to early injury.

To investigate this, we administer a controlled cortical impact (CCI) to model severe, bilateral frontal lobe injury and assess how the age at injury—either PND 17 or PND 28—impacts the development of social behavior, aggression, and frontal cortex structure. Both male and female rats were assessed longitudinally to capture changes from pre-puberty through adulthood, offering a comprehensive view of developmental trajectories post-TBI. Specifically, we aimed to examine the effects of injury age and sex on social development and hypothesized that animals injured on PND 17, especially males, would show the most pronounced social impairments. We also sought to evaluate aggression and dominance in adulthood, including potential predictors from earlier developmental periods. We hypothesized that PND 17 TBI, especially in males, would increase social aggression and dominance and early aggression would be a good predictor of this. Finally, we investigated the structural plasticity in critical social brain regions, including the OFC, mPFC, and amygdala. We predicted that PND 17 TBI would yield reduced dendritic complexity and spine density while PND 28 TBI would exhibit pronounced myelin deficits.

This multifaceted design bridges the gaps in the current literature by systematically comparing two critical developmental injury windows, assessing both sexes, and linking behavioral outcomes to underlying neural substrates. These findings provide new insight into the temporal and sex-specific vulnerabilities of the developing brain following frontal TBI and may inform age- and sex-tailored therapeutic interventions.

## Materials and Methods

### Animals

All procedures were approved by the Southern Illinois University Animal Care and Use Committee. Eighty male and female Sprague-Dawley rats (n=40/sex) (Harlan Laboratories, Indianapolis, IN) were bred in-house and maintained on a 12-h light/dark cycle with *ad libitum* access to food and water. Litters were randomized into four groups: bilateral frontal CCI (fCCI) or sham surgery on PND 17 or 28. After weaning on PND 21, rats were pair-housed with conspecifics from different groups to control for injury-related effects on play behavior (49).

### Surgical Procedures

fCCI was delivered to the mPFC on either PND 17 or 28 using a 4 mm probe (2.25 m/s velocity, 500 ms dwell, 3 mm depth), following previously validated protocols that model clinical TBI pathology (63). Sham surgeries involved anesthesia and craniotomy without impact.

### Behavioral Testing

To ensure injury did not affect motor or olfactory function, we assessed gross motor coordination using the foot fault task (FFT) on PNDs 29, 31, and 33, and olfaction (buried food and habituation/dishabituation tasks) on PND 30. Social behavior was evaluated via play behavior observations (PNDs 25, 42, 56), three-chamber task (PNDs 36-37, 43-44, 57-58), social dominance tube task (PNDs 59-60), and resident/intruder aggression task (PNDs 61-62) (Figure 1).

**Figure 1.**
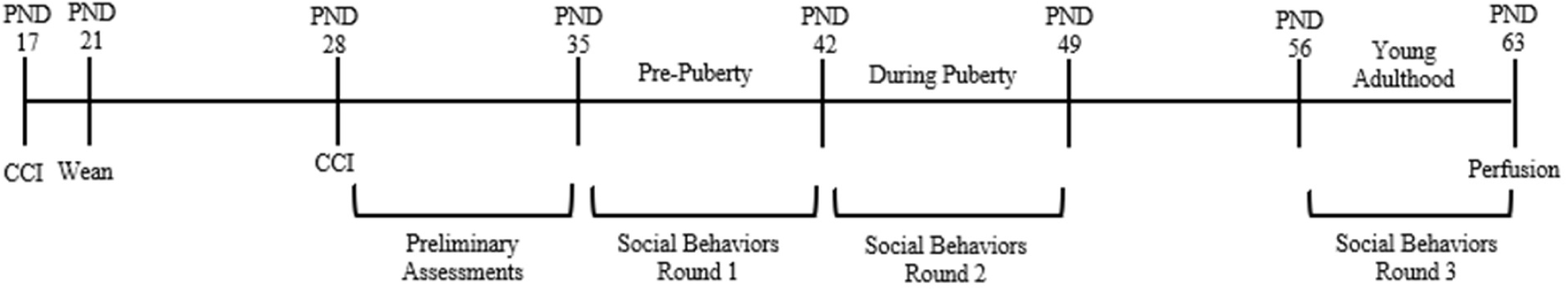
Study Procedure Timeline. *Note.* CCI, controlled cortical impact

*Preliminary Assessments.* FFT followed established methods to quantify limb coordination (63) using three trials per day to measure foot faults. Buried food and habituation/dishabituation tasks assessed basic social odor discrimination, using established protocols (64). The latency to finding the buried fruit loop was measured in the buried food task. In the habituation/dishabituation task, animals were presented an odor on a cotton applicator three times sequentially before the next odor set was presented. Odors included water, almond (Mcormick, Hunt Valley, MD; 1:100 dilution), banana (Mcormick, Hunt Valley, MD; 1:100 dilution), and two novel rat cages. Time spent sniffing the applicator was measured and habituation/dishabituation scores were calculated. Habituation scores for each odor was measured as the rate of change in time sniffing the applicator between the first trial and third trial. Dishabituation scores for each odor change was measured as the rate of change in time sniffing the new odor in the first trial versus the third trial of the previous odor. Repeated-measure ANOVAs confirmed consistent habituation scores (*F*(3.576,282.502)=1.182, *p*=0.319, η^2^=0.015) and variation in dishabituation scores (*F*(2.651,209.417)=5.287, *p*<0.01, η^2^=0.063). Thus, habituation scores were collapsed into one score, but dishabituation scores were analyzed separately.

*Play behavior observations.* Dyadic play between cage-mates was recorded and coded using previously established protocols (49,65,66). Attack and non-response behaviors were measured and served as proxies for social initiation and withdrawal.

*Three chamber task.* Social preference (novel rat vs. novel object) and memory (novel vs familiar rat) were assessed using previously describe procedures (63). Novel rats were age- and sex-matched. Time spent interacting with each stimulus (nose-touching, sniffing, or climbing of the barrier) across the ten-minute trial was used to compute preference and memory ratios (time interacting with novel rat versus novel object/familiar rat).

*Tube task.* To assess social dominance, rats were matched with a novel age- and sex-matched peer and the task was conducted as previously described (63,67). The tube was sized such that animals could not pass each other (6 cm diameter for females, 10 cm for males). Three pre-test trials established home-cage dominance, which was controlled in statistical analysis.

*Resident/intruder task.* Rats alternated as resident and intruder with novel sex- and age-matched conspecifics as previously described (68). Behaviors were recorded for 10 minutes and assessed for offensive behaviors (summation of lateral threat, upright posture, clinch attack, keep down and chase behaviors) and violence (inverse latency to initial clinch attack, calculated by subtracting the latency from the total trial duration). No trials were terminated due to excessive violence (bites to vulnerable body parts).

*Behavior coding and analysis.* Behaviors were coded using DeepLabCut (2.3.0), Behavioral Observation Research Interactive Software (BORIS, 7.13), and Simulated Behavioral Analysis (SimBA, 1.55) software. DeepLabCut was used for pose estimation (69,70), BORIS was used for manual annotations (71), and SimBA performed automated behavior classification (72). Model training parameters are provided in supplementary materials.

### Transcardial Perfusion

Animals were euthanized on PND 63 and transcardially perfused. Brains were collected and divided for either Luxol-fast blue (LFB; n=5/group/sex) or Golgi-Cox staining (n=5/group/sex). LFB tissue was post-fixed using 4% paraformaldehyde in PBS, and Golgi-Cox tissue was placed in Golgi-Cox solution (73).

### Golgi-Cox Staining

Tissue was stained based on previously established protocols (73) and complete staining procedures are outlined in supplementary material. 200 μm sections we sliced on a vibratome, and 2 coronal sections were collected of the mPFC and OFC (+3.20 mm to +2.70 mm to bregma). Pyramidal neurons in layer V of mPFC and OFC were imaged using an Olympus CX31 microscope to quantify dendritic structure (n=2/animal; total n=10/group/sex) and spine density (n=2/animal; total n=10/group/sex). Sections were images at 20x for dendritic structure analysis and 100x magnification for spine density analysis with enough images taken at different focal distances to construct an accurate and traceable z-stack. The Sholl analysis critical value, total dendrite length, and spine density were assessed using Image J (NIH, Bethesda, MD) and the SNT plugin (74).

### Luxol-Fast Blue Staining

Complete staining procedures are outlined in supplementary material. Myelin density was assessed in the mPFC and OFC (+3.70 mm to +2.70 mm relative to bregma), as well as the ventral amygdalofugal pathway and BLA (−1.80 mm to −3.14 mm relative to bregma) at 40x magnification (n=2 slices/animal/region; n=2 images/region/animal; total n=10 images/region/group/sex). Percentage of myelinated area was quantified from binary-converted images using ImageJ.

### Statistical Analysis

Sham groups were collapsed after confirming no significant differences (Table S1). Analyses were conducted in RStudio (V4.4.1). Non-parametric repeated measures ANOVA (nparLD) (75) was used for FFT results, MANOVA (npmv) (76) for olfaction, GLM (mcglm) (77) for social behaviors, and resampling-based MANOVA (MANOVA.RM) (78) for histology, with p<0.05 considered significant for all analyses.

## Results

### Preliminary Assessments

The FFT showed a significant effect of group on testing day 3 (ANOVA-Type Statistic (ATS)(1.892,56.551)=25.592, *p*<0.001, η^2^=0.461; Figure 2). *Post hoc* tests showed significantly more foot faults in PND 28 TBI versus sham animals (*p*<0.01, *d*=0.913). Suggesting that TBI on PND 28 resulted in motor deficits. To determine if these deficits were transient, *post hoc* tests between testing day 1 and 3 were compared. PND 28 TBI animals significantly reduced foot faults across days (*p*<0.01, *d*=2.011). This improvement suggests that either diaschisis resolved (79) or compensatory mechanisms were developed (80). Thus, motor deficits were likely elevated because these animals were injured temporally closer to FFT testing and were subsequently still recovering from the acute effects of injury. Given the significant improvement in performance, we conclude that there were no long-term motor deficits following PND 28 TBI.

**Figure 2.**
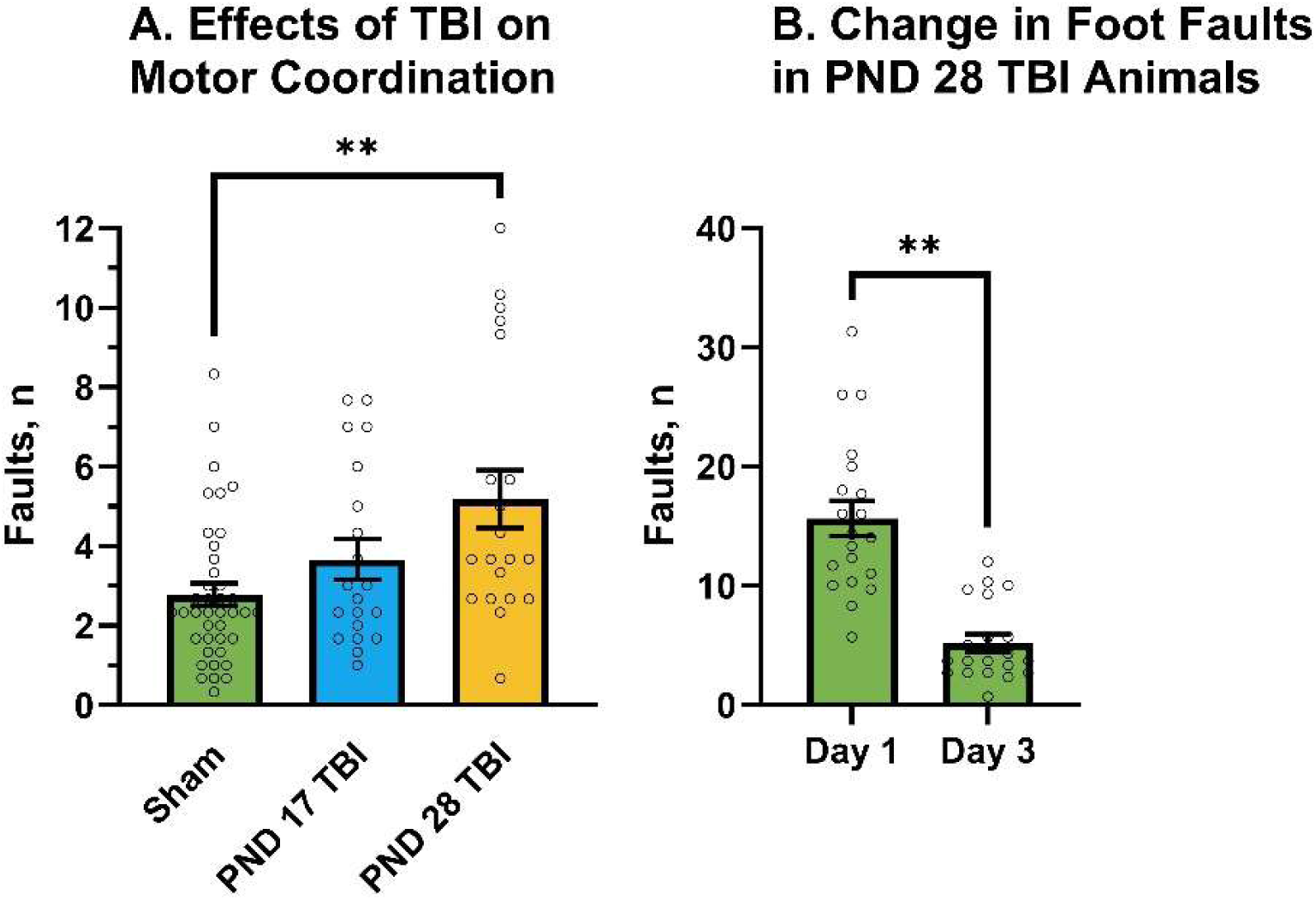
Age-Related Effects of TBI on Motor Coordination. *Note.* Graphs show group means ± SEM. A) PND 28 TBI animals displayed more foot faults on testing day three than sham animals, B) PND 28 TBI animals significantly improved motor coordination across testing days, demonstrating fewer foot faults on testing day 3 versus day 1. **, *p*<0.01

Olfactory assessments did not display significant differences between groups (ATS(8.905,306.137)=1.385, *p*=0.78, η^2^=0.039; Figure 3). Thus, even though olfactory deficits are common following TBI (81,82), especially frontal TBI (83,84), no deficits were observed in our study and thus do not confound our social behavior results.

**Figure 3.**
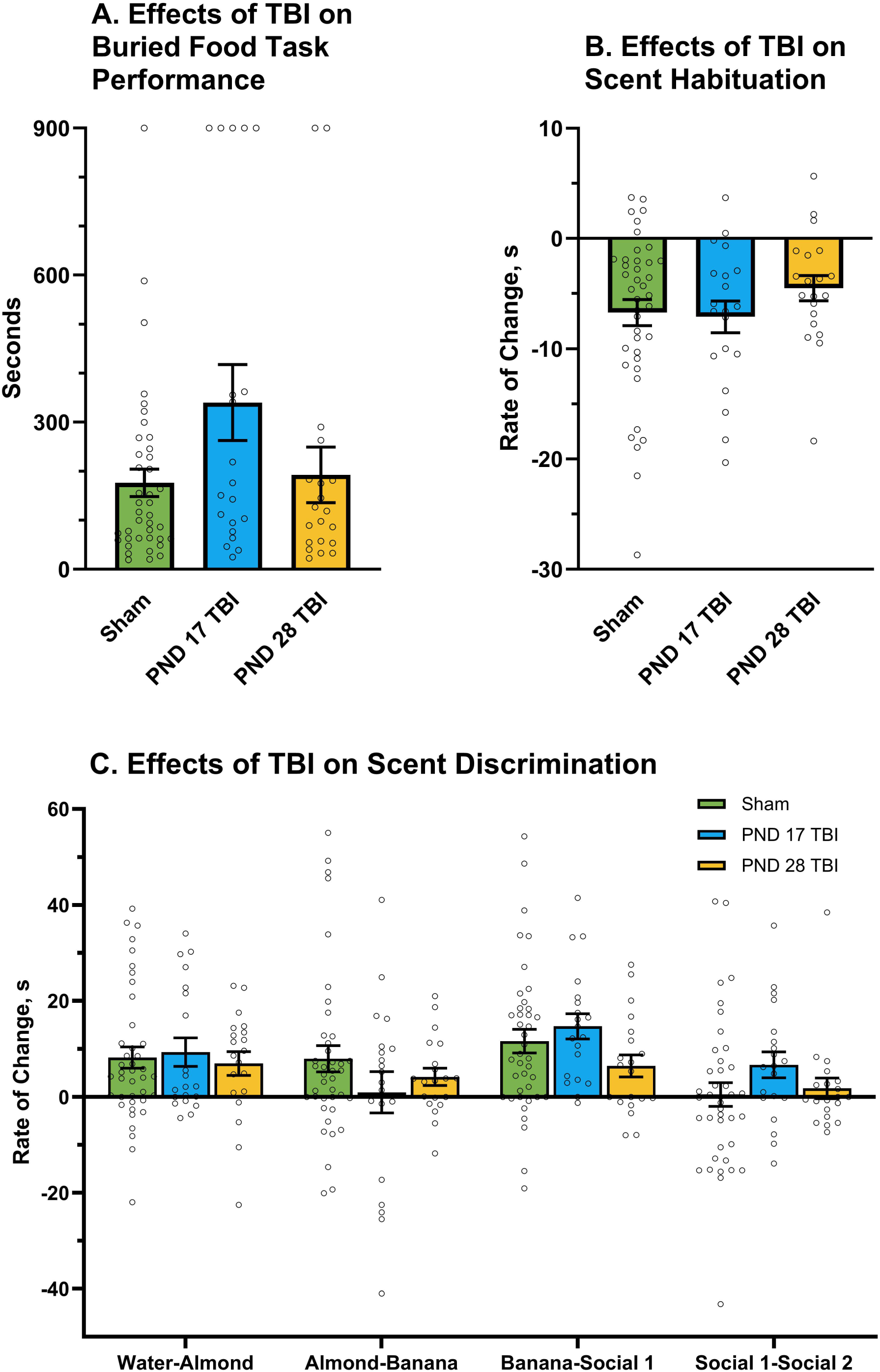
Effects of TBI on Olfaction. *Note.* Graphs show group means ± SEM. A) The latency to finding the buried fruit loop did not significantly differ between groups, B) habitation to an odor did not significantly differ between groups, C) dishabituation to an odor did not significantly differ between groups. ROC, rate of change, habituation ROC, time spent sniffing on trial 3 – time spent sniffing on trial 1 divided by two trials, dishabituation ROC, time spent sniffing on trial 1 of new odor – time spent sniffing on trial 3 of previous odor divided by two trials.

### Social Behaviors

Play initiation (Figure 4), social dominance (Figure 5), resident offensive aggression (Figure 6A and Figure 7A), and intruder offensive aggression (Figure 6B and Figure 7B) displayed significant effects of TBI. No other behaviors were significantly affected by TBI (Table 1). Graphical results of non-significant behaviors (three chamber task [Figure S1], and the impact of childhood play behavior on adult dominance [Figure S2] and violence [Figure S3]) are provided in supplementary material.

**Figure 4.**
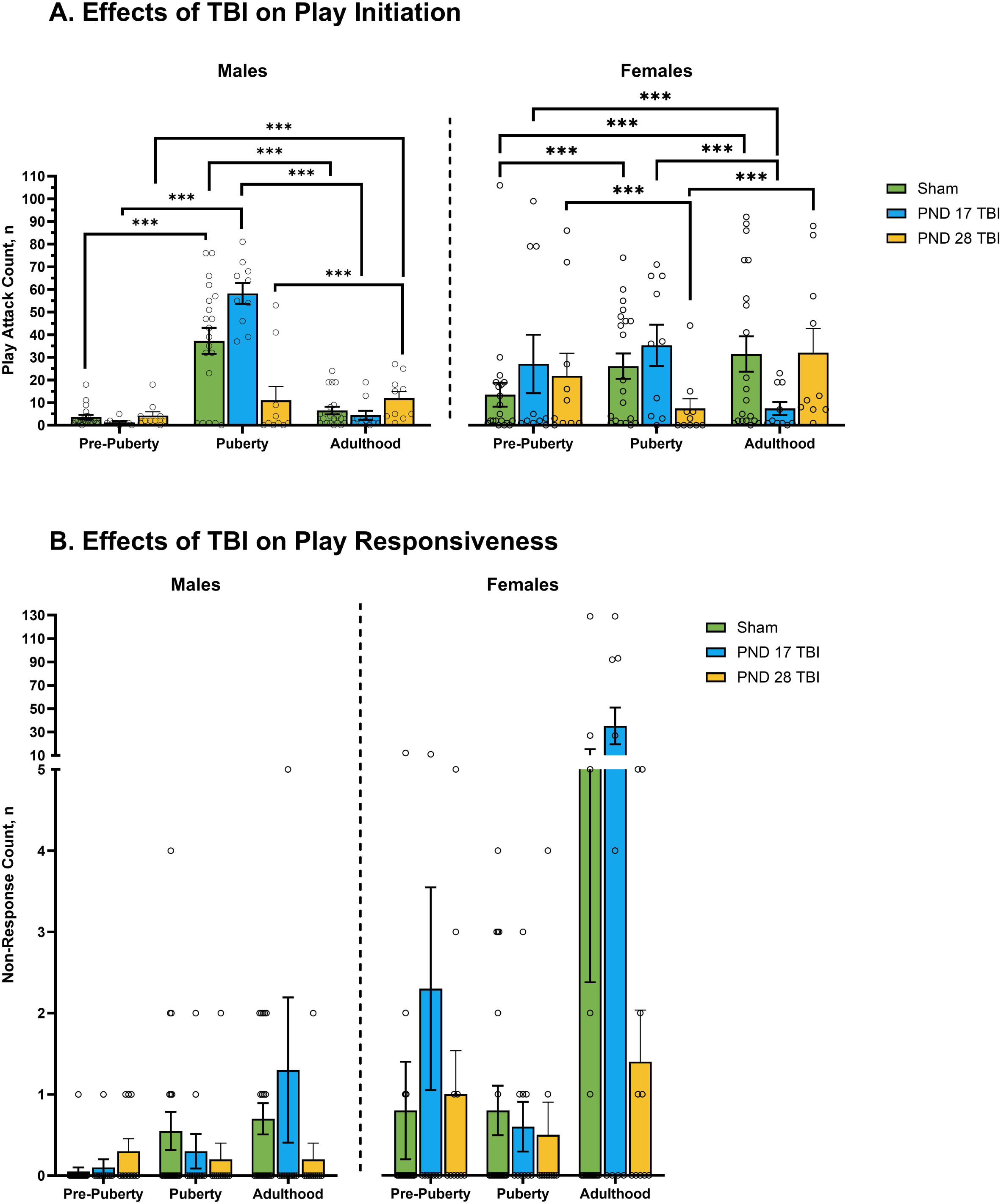
Effects of TBI on Play Behavior. *Note.* Graphs show group means ± SEM. A) TBI significantly increased play attacks compared to sham animals with PND 28 TBI animals displaying more play attacks than PND 17 TBI animals; female PND 28 TBI animals decreased play attack behavior across development. B) TBI did not significantly affect non-responsive behaviors. ***, *p*<0.001

**Figure 5.**
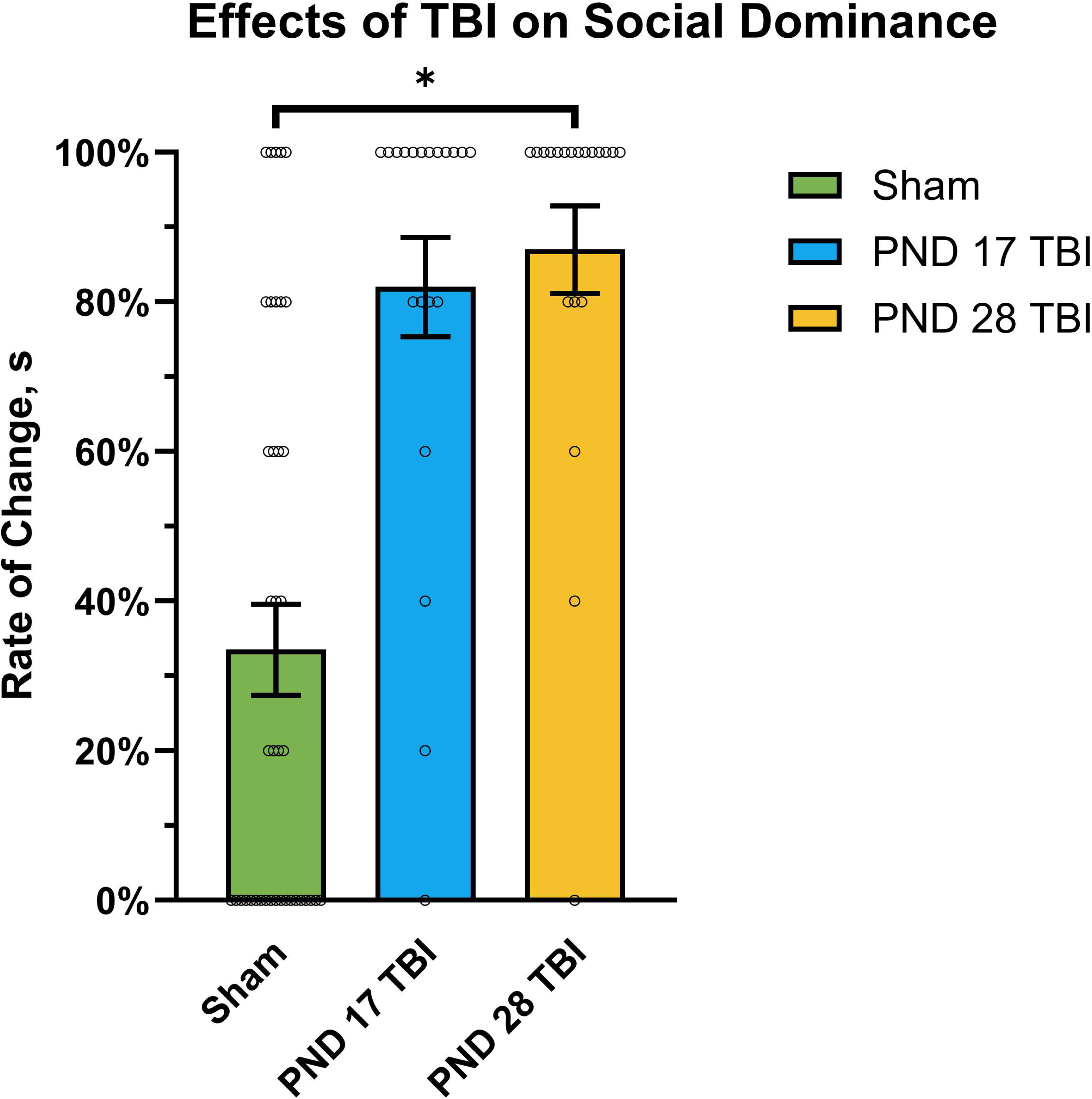
Effects of TBI on Social Dominance. *Note.* Graphs show group means ± SEM. TBI on PND 28 significantly increased dominance behaviors compared to sham animals. *, *p*<0.05.

**Figure 6.**
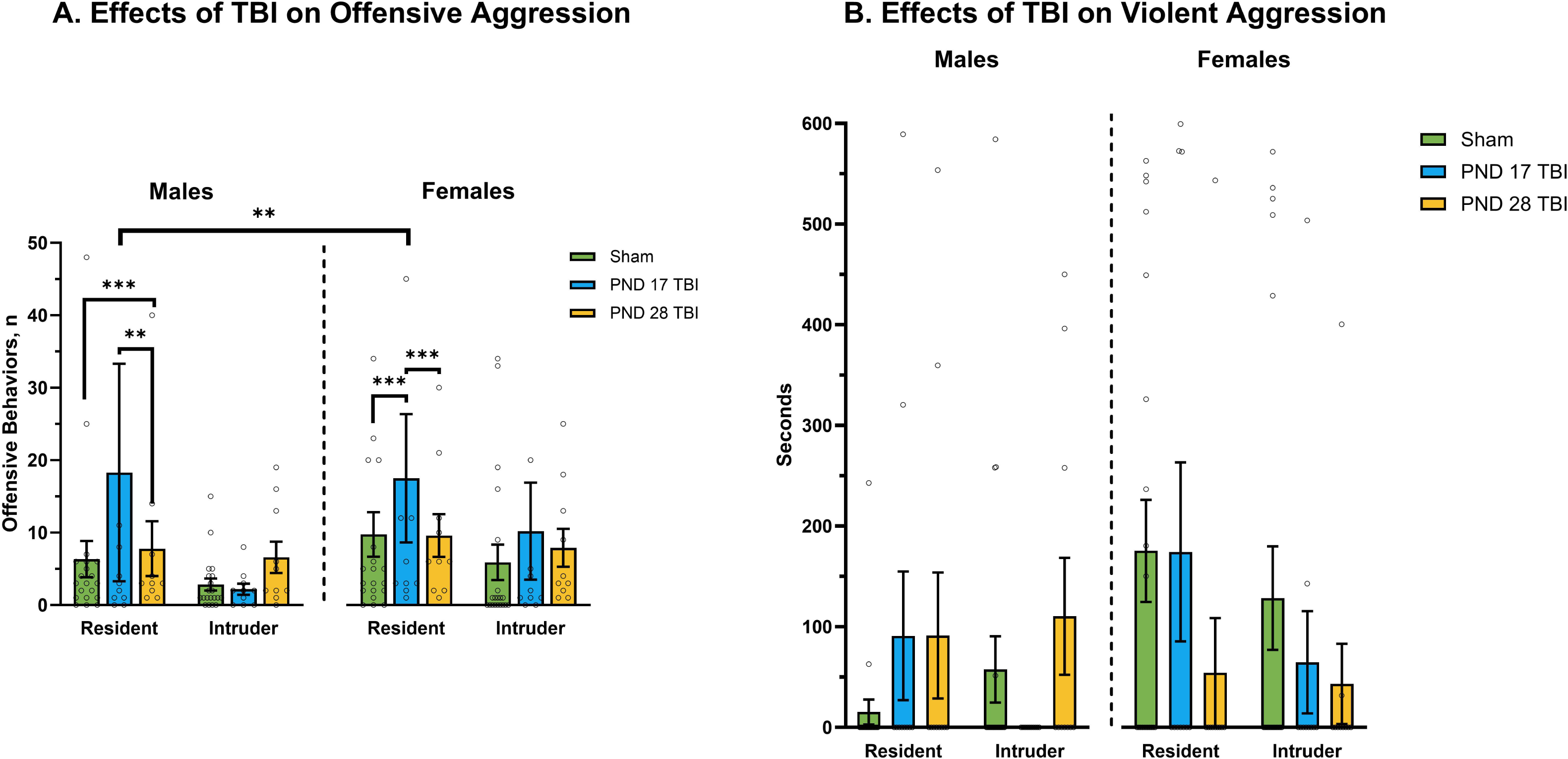
Effects of TBI on Aggression in the Resident/Intruder Task. *Note.* Graphs show group means ± SEM. A) PND 28 TBI increased resident offensive behavior in male animals while a PND 17 TBI increased resident offensive behavior in female animals, Female PND 17 TBI animals also displayed more resident offensive behaviors than male animals; there were no significant effects of sex or TBI on intruder offensive behaviors. B) There were no significant effects of TBI or sex on resident or intruder violence. **, *p*<0.01, ***, *p*<0.001.

**Figure 7.**
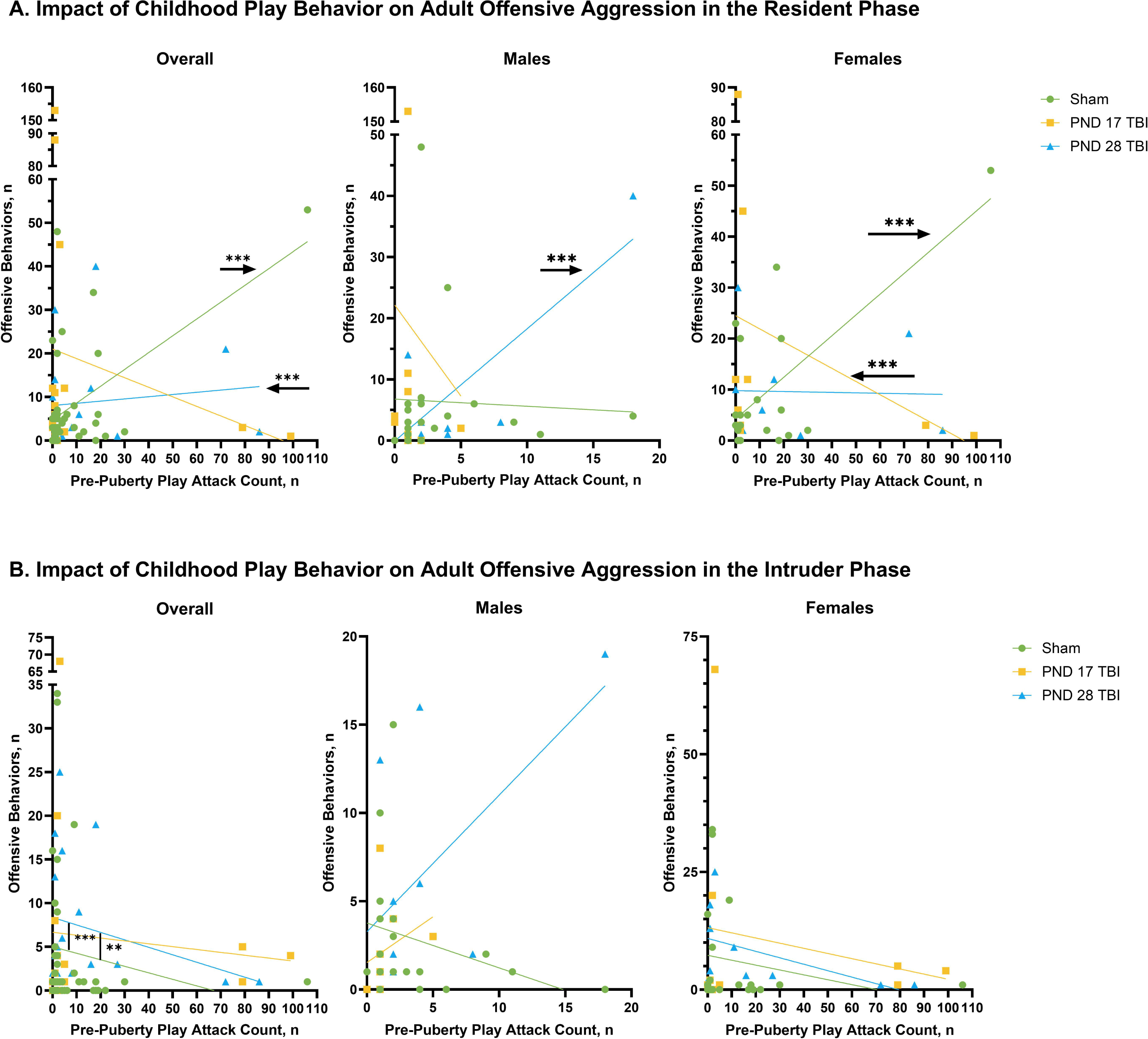
Impact of Childhood Play Behavior on Adult Offensive Aggression in the Resident/Intruder Task. *Note.* Graphs show individual data points with linear regression lines. A) Male PND 28 TBI animals displayed a positive correlation between childhood aggression and resident offensive behavior, female sham animals also displayed a positive correlation but female PND 17 TBI animals displayed a negative correlation. B) Sex did not impact the relationship between childhood play behavior and adult intruder offensive aggression but TBI on PND 28 significantly increased intruder offensive aggression when childhood play attack behavior was at or above mean levels. **, *p*<0.01, ***, *p*<0.001

**Table 1.**
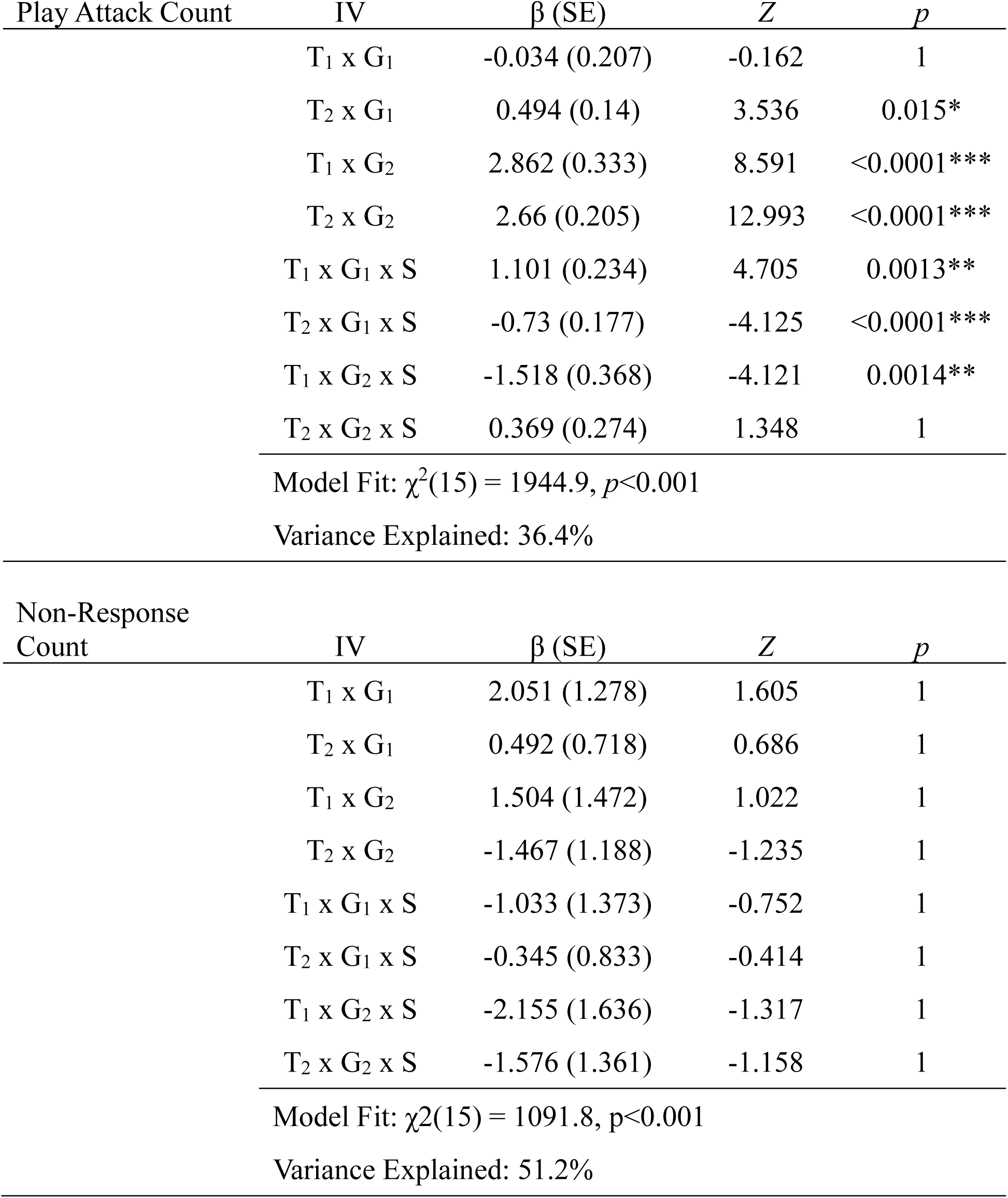
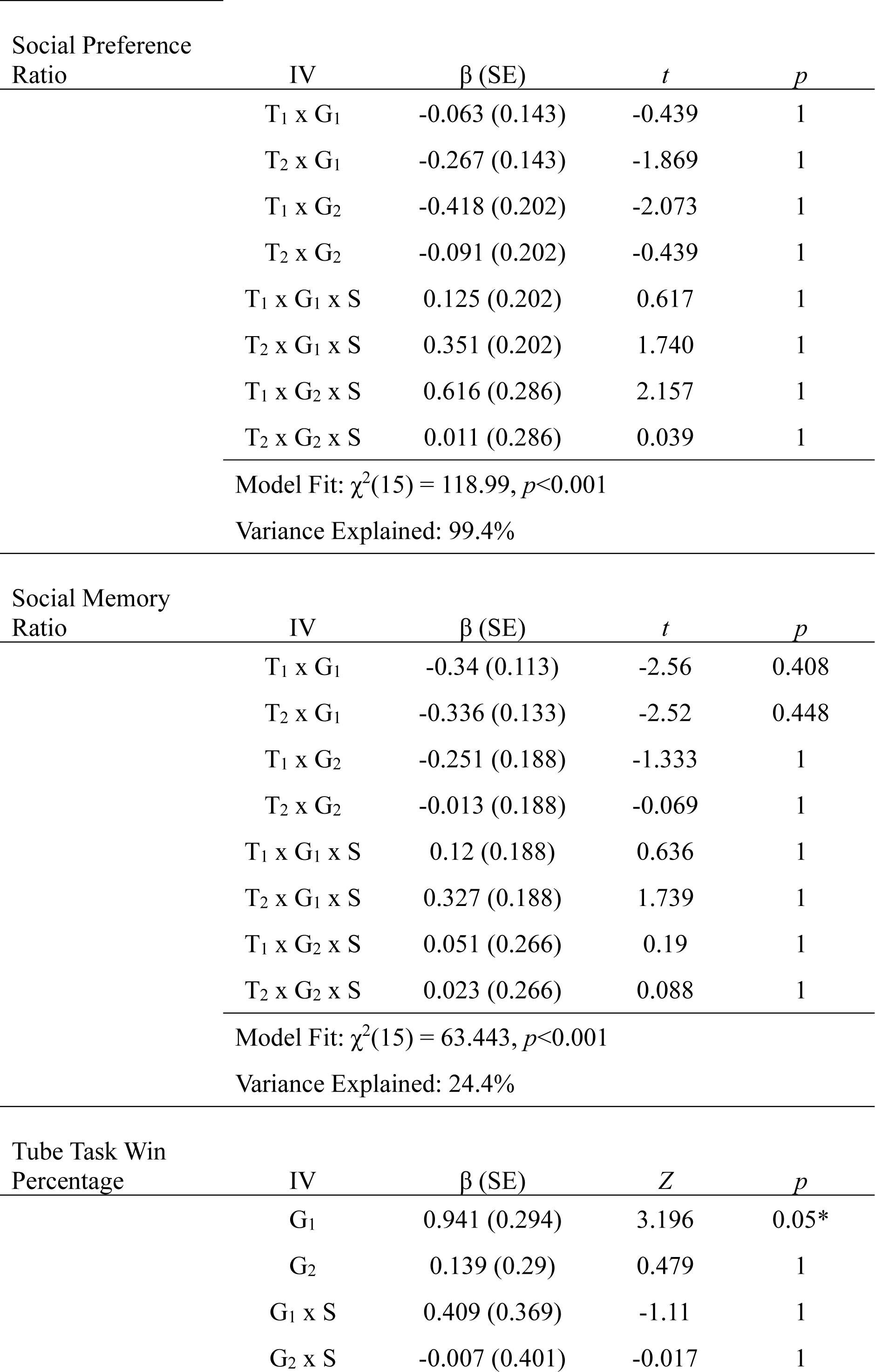
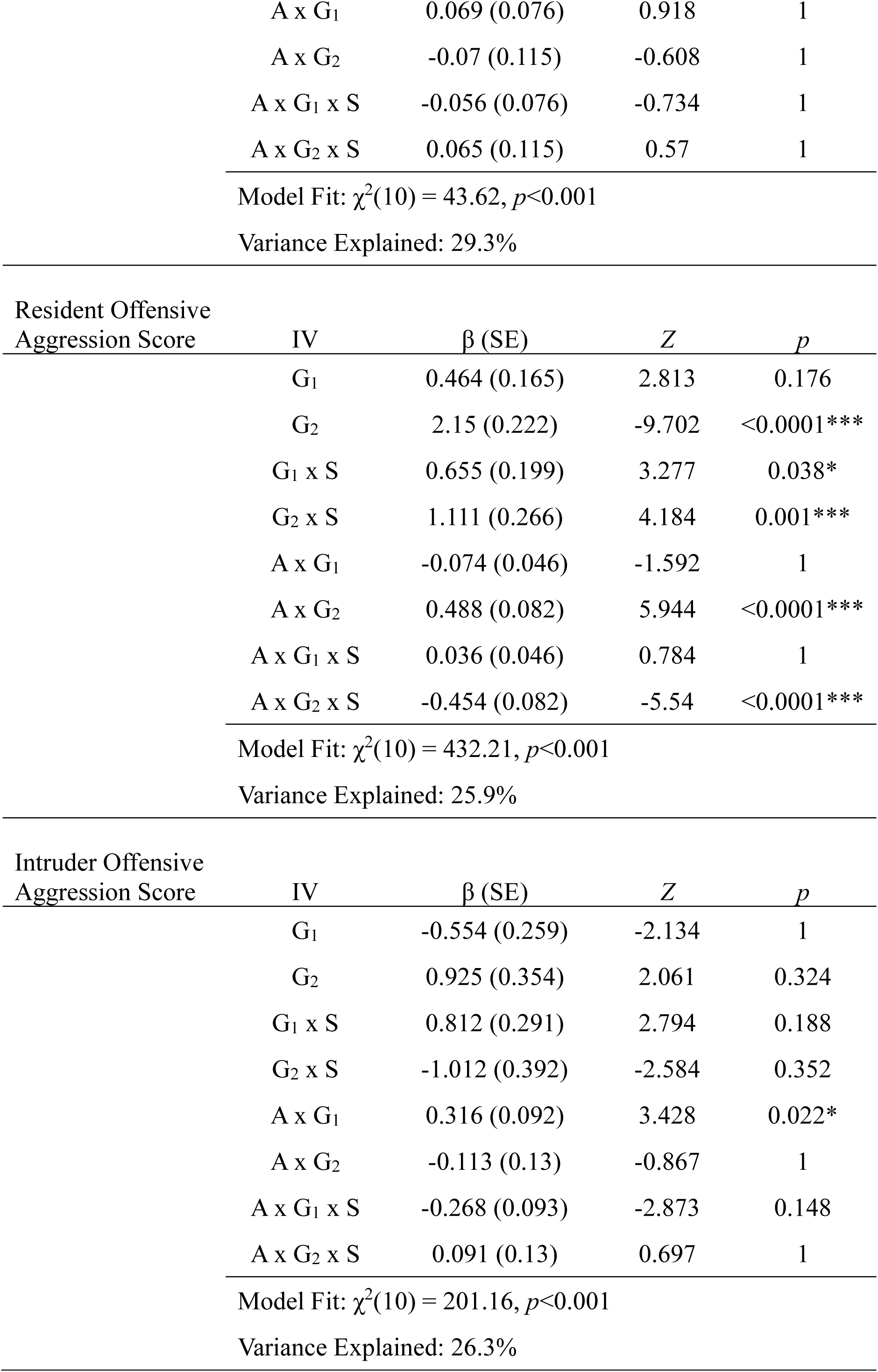
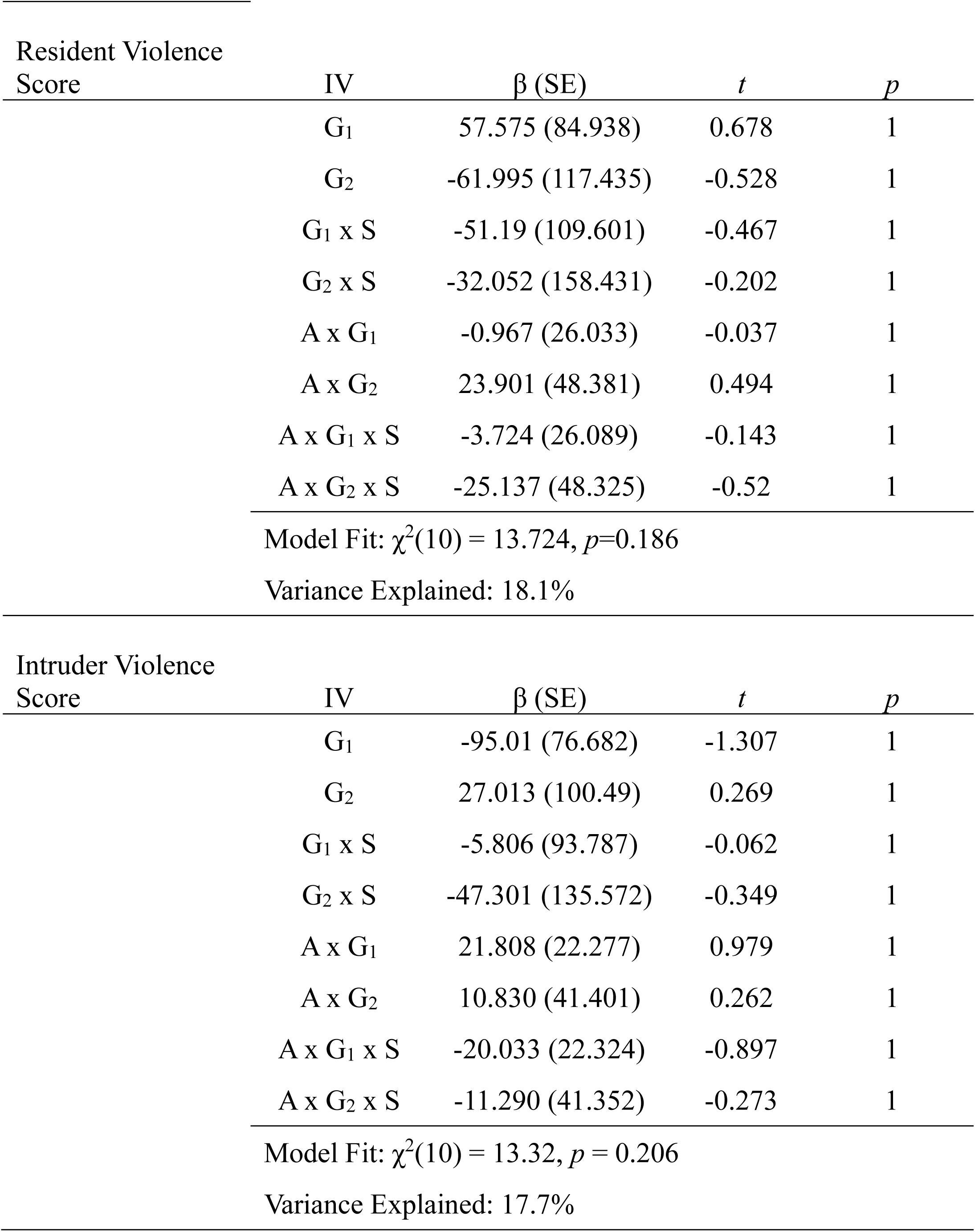
Summary of GLM Results for Social Behaviors. *Note.* A, number of play attacks at the pre-puberty time point, G_1_, sham group compared to both TBI groups, G_2_, PND 17 TBI group compared to the PND 28 TBI group, T_1_, pre-puberty time point compared to puberty time point, T_2_, adult time point compared to puberty time point, S, sex, *, *p*<0.05, **, *p*<0.01, ***, *p*<0.001

| Play Attack Count | IV | $\beta$ (SE) | Z | p |
| --- | --- | --- | --- | --- |
| T<sub>1</sub> x G<sub>1</sub> |  | -0.034 (0.207) | -0.162 | 1 |
| T<sub>2</sub> x G<sub>1</sub> |  | 0.494 (0.14) | 3.536 | 0.015* |
| T<sub>1</sub> x G<sub>2</sub> |  | 2.862 (0.333) | 8.591 | <0.0001*** |
| T<sub>2</sub> x G<sub>2</sub> |  | 2.66 (0.205) | 12.993 | <0.0001*** |
| T<sub>1</sub> x G<sub>1</sub> x S |  | 1.101 (0.234) | 4.705 | 0.0013** |
| T<sub>2</sub> x G<sub>1</sub> x S |  | -0.73 (0.177) | -4.125 | <0.0001*** |
| T<sub>1</sub> x G<sub>2</sub> x S |  | -1.518 (0.368) | -4.121 | 0.0014** |
| T<sub>2</sub> x G<sub>2</sub> x S |  | 0.369 (0.274) | 1.348 | 1 |
| Model Fit: $\chi^2(15) = 1944.9, p < 0.001$ | | | | |
| Variance Explained: 36.4% |  |  |  |  |
| Non-Response Count | IV | $\beta$ (SE) | Z | p |
| T<sub>1</sub> x G<sub>1</sub> |  | 2.051 (1.278) | 1.605 | 1 |
| T<sub>2</sub> x G<sub>1</sub> |  | 0.492 (0.718) | 0.686 | 1 |
| T<sub>1</sub> x G<sub>2</sub> |  | 1.504 (1.472) | 1.022 | 1 |
| T<sub>2</sub> x G<sub>2</sub> |  | -1.467 (1.188) | -1.235 | 1 |
| T<sub>1</sub> x G<sub>1</sub> x S |  | -1.033 (1.373) | -0.752 | 1 |
| T<sub>2</sub> x G<sub>1</sub> x S |  | -0.345 (0.833) | -0.414 | 1 |
| T<sub>1</sub> x G<sub>2</sub> x S |  | -2.155 (1.636) | -1.317 | 1 |
| T<sub>2</sub> x G<sub>2</sub> x S |  | -1.576 (1.361) | -1.158 | 1 |
| Model Fit: $\chi^2(15) = 1091.8, p < 0.001$ | | | | |
| Variance Explained: 51.2% |  |  |  |  |

| Social Preference Ratio | IV | $\beta$ (SE) | $t$ | $p$ |
| --- | --- | --- | --- | --- |
|  | T<sub>1</sub> x G<sub>1</sub> | -0.063 (0.143) | -0.439 | 1 |
|  | T<sub>2</sub> x G<sub>1</sub> | -0.267 (0.143) | -1.869 | 1 |
|  | T<sub>1</sub> x G<sub>2</sub> | -0.418 (0.202) | -2.073 | 1 |
|  | T<sub>2</sub> x G<sub>2</sub> | -0.091 (0.202) | -0.439 | 1 |
|  | T<sub>1</sub> x G<sub>1</sub> x S | 0.125 (0.202) | 0.617 | 1 |
|  | T<sub>2</sub> x G<sub>1</sub> x S | 0.351 (0.202) | 1.740 | 1 |
|  | T<sub>1</sub> x G<sub>2</sub> x S | 0.616 (0.286) | 2.157 | 1 |
|  | T<sub>2</sub> x G<sub>2</sub> x S | 0.011 (0.286) | 0.039 | 1 |
| Model Fit: $\chi^2(15) = 118.99, p < 0.001$ | | | | |
| Variance Explained: 99.4% |  |  |  |  |

| Social Memory Ratio | IV | $\beta$ (SE) | $t$ | $p$ |
| --- | --- | --- | --- | --- |
|  | T<sub>1</sub> x G<sub>1</sub> | -0.34 (0.113) | -2.56 | 0.408 |
|  | T<sub>2</sub> x G<sub>1</sub> | -0.336 (0.133) | -2.52 | 0.448 |
|  | T<sub>1</sub> x G<sub>2</sub> | -0.251 (0.188) | -1.333 | 1 |
|  | T<sub>2</sub> x G<sub>2</sub> | -0.013 (0.188) | -0.069 | 1 |
|  | T<sub>1</sub> x G<sub>1</sub> x S | 0.12 (0.188) | 0.636 | 1 |
|  | T<sub>2</sub> x G<sub>1</sub> x S | 0.327 (0.188) | 1.739 | 1 |
|  | T<sub>1</sub> x G<sub>2</sub> x S | 0.051 (0.266) | 0.19 | 1 |
|  | T<sub>2</sub> x G<sub>2</sub> x S | 0.023 (0.266) | 0.088 | 1 |
| Model Fit: $\chi^2(15) = 63.443, p < 0.001$ | | | | |
| Variance Explained: 24.4% |  |  |  |  |

| Tube Task Win Percentage | IV | $\beta$ (SE) | $Z$ | $p$ |
| --- | --- | --- | --- | --- |
|  | G<sub>1</sub> | 0.941 (0.294) | 3.196 | 0.05* |
|  | G<sub>2</sub> | 0.139 (0.29) | 0.479 | 1 |
|  | G<sub>1</sub> x S | 0.409 (0.369) | -1.11 | 1 |
|  | G<sub>2</sub> x S | -0.007 (0.401) | -0.017 | 1 |

|  |  |  |  |
| --- | --- | --- | --- |
| A x G<sub>1</sub> | 0.069 (0.076) | 0.918 | 1 |
| A x G<sub>2</sub> | -0.07 (0.115) | -0.608 | 1 |
| A x G<sub>1</sub> x S | -0.056 (0.076) | -0.734 | 1 |
| A x G<sub>2</sub> x S | 0.065 (0.115) | 0.57 | 1 |
| Model Fit: $\chi^2(10) = 43.62, p < 0.001$ | | | |
| Variance Explained: 29.3% |  |  |  |

| Resident Offensive Aggression Score | IV | $\beta$ (SE) | Z | p |
| --- | --- | --- | --- | --- |
| G<sub>1</sub> |  | 0.464 (0.165) | 2.813 | 0.176 |
| G<sub>2</sub> |  | 2.15 (0.222) | -9.702 | <0.0001*** |
| G<sub>1</sub> x S |  | 0.655 (0.199) | 3.277 | 0.038* |
| G<sub>2</sub> x S |  | 1.111 (0.266) | 4.184 | 0.001*** |
| A x G<sub>1</sub> |  | -0.074 (0.046) | -1.592 | 1 |
| A x G<sub>2</sub> |  | 0.488 (0.082) | 5.944 | <0.0001*** |
| A x G<sub>1</sub> x S |  | 0.036 (0.046) | 0.784 | 1 |
| A x G<sub>2</sub> x S |  | -0.454 (0.082) | -5.54 | <0.0001*** |
| Model Fit: $\chi^2(10) = 432.21, p < 0.001$ | | | | |
| Variance Explained: 25.9% |  |  |  |  |

| Intruder Offensive Aggression Score | IV | $\beta$ (SE) | Z | p |
| --- | --- | --- | --- | --- |
| G<sub>1</sub> |  | -0.554 (0.259) | -2.134 | 1 |
| G<sub>2</sub> |  | 0.925 (0.354) | 2.061 | 0.324 |
| G<sub>1</sub> x S |  | 0.812 (0.291) | 2.794 | 0.188 |
| G<sub>2</sub> x S |  | -1.012 (0.392) | -2.584 | 0.352 |
| A x G<sub>1</sub> |  | 0.316 (0.092) | 3.428 | 0.022* |
| A x G<sub>2</sub> |  | -0.113 (0.13) | -0.867 | 1 |
| A x G<sub>1</sub> x S |  | -0.268 (0.093) | -2.873 | 0.148 |
| A x G<sub>2</sub> x S |  | 0.091 (0.13) | 0.697 | 1 |
| Model Fit: $\chi^2(10) = 201.16, p < 0.001$ | | | | |
| Variance Explained: 26.3% |  |  |  |  |

| Resident Violence Score | IV | $\beta$ (SE) | $t$ | $p$ |
| --- | --- | --- | --- | --- |
|  | G<sub>1</sub> | 57.575 (84.938) | 0.678 | 1 |
|  | G<sub>2</sub> | -61.995 (117.435) | -0.528 | 1 |
|  | G<sub>1</sub> x S | -51.19 (109.601) | -0.467 | 1 |
|  | G<sub>2</sub> x S | -32.052 (158.431) | -0.202 | 1 |
|  | A x G<sub>1</sub> | -0.967 (26.033) | -0.037 | 1 |
|  | A x G<sub>2</sub> | 23.901 (48.381) | 0.494 | 1 |
|  | A x G<sub>1</sub> x S | -3.724 (26.089) | -0.143 | 1 |
|  | A x G<sub>2</sub> x S | -25.137 (48.325) | -0.52 | 1 |
| Model Fit: $\chi^2(10) = 13.724, p=0.186$ | | | | |
| Variance Explained: 18.1% |  |  |  |  |
| Intruder Violence Score | IV | $\beta$ (SE) | $t$ | $p$ |
|  | G<sub>1</sub> | -95.01 (76.682) | -1.307 | 1 |
|  | G<sub>2</sub> | 27.013 (100.49) | 0.269 | 1 |
|  | G<sub>1</sub> x S | -5.806 (93.787) | -0.062 | 1 |
|  | G<sub>2</sub> x S | -47.301 (135.572) | -0.349 | 1 |
|  | A x G<sub>1</sub> | 21.808 (22.277) | 0.979 | 1 |
|  | A x G<sub>2</sub> | 10.830 (41.401) | 0.262 | 1 |
|  | A x G<sub>1</sub> x S | -20.033 (22.324) | -0.897 | 1 |
|  | A x G<sub>2</sub> x S | -11.290 (41.352) | -0.273 | 1 |
| Model Fit: $\chi^2(10) = 13.32, p = 0.206$ | | | | |
| Variance Explained: 17.7% |  |  |  |  |

*Play initiation.* Injury status (*p*<0.05) and age at injury (*p*<0.001) significantly affected play initiation behaviors, as measured by play attack count, between puberty and adulthood (Table 1). Sex further moderated this relationship, imparting significant affects on play attack levels based on injury status between pre-puberty and puberty (*p*<0.01), and puberty and adulthood (*p*<0.001), as well as based on age at injury between pre-puberty and puberty (*p*<0.01) (Table 1, Figure 4A).

*Post hoc* analyses displayed significant effects across development and groups. Across development, female sham animals increased play attacks into puberty (*p*<0.001, *d*=0.518) and maintained that level into adulthood. Meanwhile, male sham animals displayed an inverted U-shaped pattern with increased play attacks leading into puberty (p<0.001, *d*=1.831) and then decreasing play attacks into adulthood (*p*<0.001, *d*=1.629) back to pre-puberty levels. These results signify that there are sex-specific differences in play behavior across development. Meanwhile, female PND 17 TBI animals did not display any differences in play attacks until after puberty where play attacks decreased into adulthood compared to both pre-puberty (*p*<0.001, d=0.666) and puberty time points (*p*<0.001, *d*=1.303). While male PND 17 TBI animals displayed the same developmental pattern as male sham animals, by increasing play attacks into puberty (*p*<0.001, *d*=5.518) then decreasing into adulthood (*p*<0.001, *d*=4.803) back to pre-puberty levels. Finally, female PND 28 TBI animals displayed a U-shaped pattern, decreasing play attacks into puberty (*p*<0.001, *d*=0.592) then increasing into adulthood (*p*<0.001, *d*=0.96) back to pre-puberty levels. However, male PND 28 TBI animals increased play attacks into puberty (*p*<0.001, *d*=0.472) and remained stable into adulthood. Thus, male PND 28 TBI animals displayed a similar pattern as female sham animals. Overall, TBI caused age- and sex-specific play initiation deficits whereby males injured on PND 17 and females injured on PND 28 exhibited the most deficits across development.

Across groups at each time point, there were sex-specific effects at each developmental time point. At the pre-puberty time point, while play attack levels in male animals across groups did not significantly differ, both PND 28 (*p*<0.001, *d*=0.408) and PND 17 TBI (*p*<0.001, *d*=0.298) increased play attacks in females but there were no differences based on age of injury. During puberty, male animals displayed an age-specific effect of TBI with increased play attacks observed in PND 17 TBI animals compared to both sham (*p*<0.001, *d*=0.49) and PND 28 TBI animals (*p*<0.001, *d*=2.757), and decreased play attacks observed in PND 28 TBI animals compared to sham animals (*p*<0.001, *d*=1.156). Meanwhile, in female animals, only the PND 28 TBI decreased play attacks compared to both sham (*p*<0.001, *d*=0.926) and PND 17 TBI animals (*p*<0.001, *d*=1.235). Finally, in adulthood, only the PND 28 TBI in males decreased play attacks compared to both sham (*p*<0.01, *d*=0.629) and PND 17 TBI animals (*p*<0.001, *d*=0.629). Conversely, only the PND 17 TBI in females decreased play attacks in adulthood compared to both sham (*p*<0.001, *d*=0.945) and PND 28 TBI animals (*p*=0.0125, *d*=0.945). Thus, TBI imparts an age- and sex-specific effect on the development of play behavior. Males with a PND 17 TBI retain higher levels of functioning throughout adulthood, whereas females with a PND 28 TBI display a suppression of play initiation.

Finally, significant sex differences were observed across development. In sham animals, females displayed more play attacks compared to males at the pre-puberty (*p*<0.001, *d*=0.583) and adult time points (*p*<0.001, *d*=0.991), and fewer play attacks during puberty (*p*<0.001, *d*=0.44). Thus, female rats display higher levels of play initiation than males except for at puberty. In PND 17 TBI animals, females also displayed more play attacks than males at the pre-puberty time point (*p*<0.001, *d*=0.894) and fewer play attacks during puberty (*p*<00.001, *d*=1.003) but there were no sex differences in adulthood. Finally, in PND 28 TBI animals, females matched sham patterns by displaying more play attacks prior to puberty (*p*<0.001, *d*=0.774) and in adulthood (*p*<0.001, *d*=0.816) but there were no sex differences during puberty. These results further display a sex- and age-specific effect of TBI where females, but not males, display increases in play initiation soon after the injury but injury effects evolve across development such that a PND 17 TBI in females decreased play initiation and a PND 28 TBI in males increased play initiation.

*Tube Task.* In the tube task, TBI significantly increased social dominance behaviors (*p*=0.05), but age of injury, sex, or childhood play attack levels did not moderate this relationship (Figure 5, Table 1). *Post hoc* analysis found that PND 28 TBI animals displayed more wins in the tube task than sham animals (*p*<0.05, *d*=1.624), but PND 17 TBI animals did not differ from PND 28 (*p* = 1, *d* = 0.179) or sham animals (*p* = 0.536, *d* = 1.41). Thus, a TBI on PND 28 increased social dominance but one on PND 17 did not. However, the effect size for PND 17 TBI animals was substantial suggesting that our sample size was insufficient to detect differences.

*Resident offensive aggression.* While TBI did not significantly impact resident offensive aggression scores, that age at injury significantly affected resident offensive aggression (*p*<0.001) (Figure 6A, Table 1). Sex moderated this relationship for both injury status (*p*<0.05) and age at injury (*p*<0.001). Finally, childhood play attack levels predicted adult resident aggression based on age at injury (*p*<0.001) and sex moderated this effect (*p*<0.001) (Figure 7A, Table 1).

*Post hoc* analyses demonstrated an age- and sex-specific effect on resident offensive aggression. In male animals, only a PND 28 TBI increased resident offensive behaviors compared to sham (p<0.001, *d*=0.125) and PND 17 TBI animals (*p*<0.01, *d*=0.303). Conversely, only a PND 17 TBI increased resident offensive aggression in female animals compared to sham (*p*<0.001, *d*=0.351) and PND 28 TBI animals (*p*<0.001, *d*=0.379). Across sexes, only female PND 17 TBI animals displayed more offensive aggression than male animals (*p*<0.01, *d*=0.021). Finally, predictability of pre-puberty play attacks on adult resident offensive aggression displayed a sex- and age-specific effect. While sham males did not display a significant relationship between pre-puberty play attacks and adult resident offensive aggression, female sham animals displayed a positive correlation (*p*<0.001, *d*=0.542), suggesting that aggression remains consistent across development but only in females. However, TBI moderated this relationship in an age-specific manner. A PND 17 TBI resulted in a negative correlation in female animals (*p*<0.001, *d*=0.7) but had no effect on males. While a PND 28 TBI resulted in a positive correlation for male animals (*p*<0.001, *d*=3.37) but had no effect on females. Once again, a PND 17 TBI in females and PND 28 TBI in males caused the largest effect on behavior but in opposite directions. Our results indicate that PND 17 TBI in females inverts the normal development of aggression while a PND 28 TBI in males increases aggression consistently across development.

*Intruder offensive aggression.* TBI and sex did not impact intruder offensive aggression (Figure 6B, Table 1), however pre-puberty play attacks did significantly predict adult intruder offensive aggression based on injury status (*p*<0.05) (Figure 7B, Table 1). *Post hoc* tests show that PND 28 TBI increased intruder aggression but only at mean (*p*<0.001, *d*=1.933) or high (i.e. +1 standard deviation) levels (*p*<0.01, *d*=5.434) of play attacks in childhood. This result further clarifies the effects of a PND 28 TBI on aggression, whereby aggression will only be increased in adulthood if elevated aggressive was also displayed in childhood. However, this effect is only observed when intruding on another animal’s territory instead of protecting their home territory.

### Histology

While resampling-based MANOVA analysis did not find any significant effects of interest, there were several large effect sizes that indicate the potential for significant effects if sample sizes were larger (Table 2). Thus, we are currently collecting larger sample sizes in order to reevaluate our results. However, large effect sizes (i.e. η^2^≥0.14) were observed for dendrite length in basal OFC dendrites (*p*=0.031, η^2^=0.17), spine density in the mPFC (*p*=0.036, η^2^=0.17), and myelin ratio in the OFC (*p*=0.013, η^2^=0.21) and mPFC (*p*=0.063, η^2^=0.14) based on group status but not sex. This implies that significant results would have been found if sample sizes were higher and the data trends we observed will be discussed.

**Table 2.** Summary of Resampling MANOVA Results for Histology Analysis. *Note.* Degrees of freedom for all analyses had a numerator of 2, and a denominator of 78. AFP, amygdalofugal pathway, BLA, basolateral amygdala, MATS, modified ANOVA-type statistic, mPFC, medial prefrontal cortex, OFC, orbitofrontal cortex. #, large effect size.

| Critical Value |  |  | MATS | *p* | η<sup>2</sup> |
| --- | --- | --- | --- | --- | --- |
| OFC | Apical | Group | 2.474 | 0.325 | 0.06 |
|  |  | Group*Sex | 0.585 | 0.585 | 0.03 |
|  | Basal | Group | 4.780 | 0.115 | 0.11 |
|  |  | Group*Sex | 4.22 | 0.135 | 0.1 |
| mPFC | Apical | Group | 1.519 | 0.494 | 0.04 |
|  |  | Group*Sex | 2.582 | 0.268 | 0.03 |
|  | Basal | Group | 1.349 | 0.524 | 0.03 |
|  |  | Group*Sex | 2.501 | 0.272 | 0.06 |
| Dendrite Length |  |  | MATS | *p* | η<sup>2</sup> |
| OFC | Apical | Group | 1.47 | 0.501 | 0.04 |
|  |  | Group*Sex | 2.199 | 0.345 | 0.05 |
|  | Basal | Group | 7.924 | 0.031 | 0.17# |
|  |  | Group*Sex | 6.05 | 0.069 | 0.13 |
| mPFC | Apical | Group | 1.898 | 0.411 | 0.05 |
|  |  | Group*Sex | 2.582 | 0.268 | 0.06 |
|  | Basal | Group | 2.476 | 0.312 | 0.06 |
|  |  | Group*Sex | 1.888 | 0.953 | 0.05 |
| Spine Density |  |  | MATS | *p* | η<sup>2</sup> |
| OFC |  | Group | 0.781 | 0.68 | 0.02 |
|  |  | Group*Sex | 0.384 | 0.85 | 0.01 |
| mPFC |  | Group | 7.983 | 0.036 | 0.17# |
|  |  | Group*Sex | 0.403 | 0.831 | 0.01 |

| Myelin Ratio | | MATS | *p* | $\eta^2$ |
| --- | --- | --- | --- | --- |
| OFC | Group | 10.473 | 0.013 | 0.21# |
|  | Group*Sex | 4.105 | 0.142 | 0.1 |
| mPFC | Group | 6.331 | 0.063 | 0.14# |
|  | Group*Sex | 1.207 | 0.551 | 0.03 |
| AFP | Group | 1.437 | 0.497 | 0.04 |
|  | Group*Sex | 1.874 | 0.388 | 0.05 |
| BLA | Group | 0.68 | 0.715 | 0.02 |
|  | Group*Sex | 0.8 | 0.692 | 0.02 |

*Basal OFC dendrite length.* The large effect size indicates that TBI impacted basal dendrite length in the OFC (Figure 8), but sample sizes were not large enough (n=2/animal; n=10/group/sex) to detect a significant difference. Thus, results indicate that TBI on PND 17 but not PND 28 reduced dendrite length in basal OFC neurons. This provides support for our hypothesis and indicates that TBI on PND 17 decreases local neuronal connections within the OFC.

**Figure 8.**
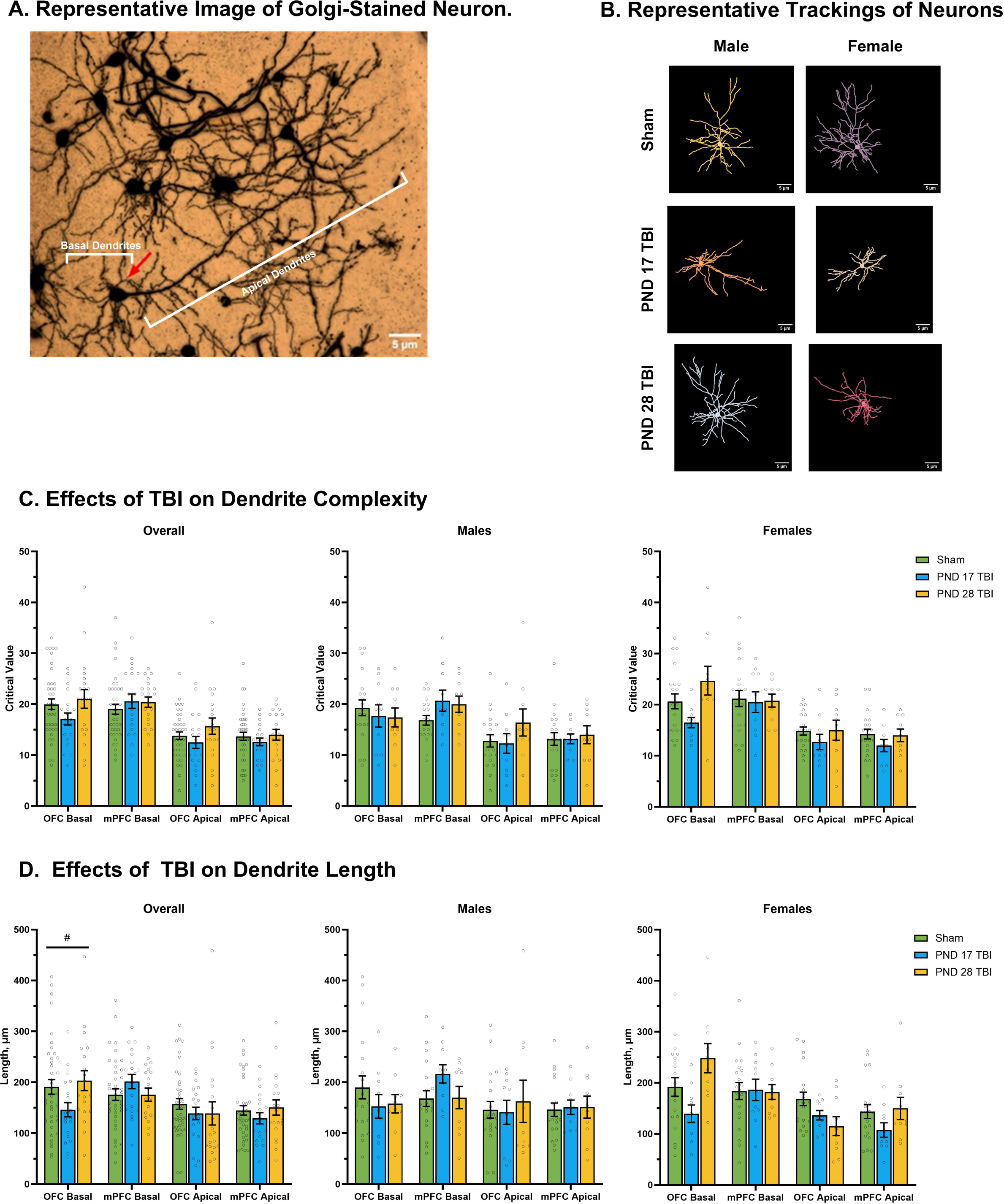
Effects of TBI on Dendrite Morphology. *Note.* Graphs show group means ± SEM. A-B) Scale bar = 5 µm. C) Neither TBI nor sex significantly impacted dendrite complexity in the OFC or mPFC. D) Neither TBI nor sex significantly impacted dendrite length in the mPFC or OFC, however, there was a large effect size on basal OFC dendrites, suggesting that TBI impacted dendrite length, but the sample size was too small. #, η^2^≥0.14

*mPFC spine density.* TBI also likely impacts spine density in the mPFC (Figure 9), but sample sizes were not large enough to detect a significant difference. Based on our results, TBI decreased spine density regardless of injury day. This partially supports our hypothesis where we expected decreases in PND 17 TBI animals but PND 28 TBI animals also trended towards spine density deficits. This suggests that TBI to the mPFC, regardless of injury day, decreased synaptic density within the mPFC.

**Figure 9.**
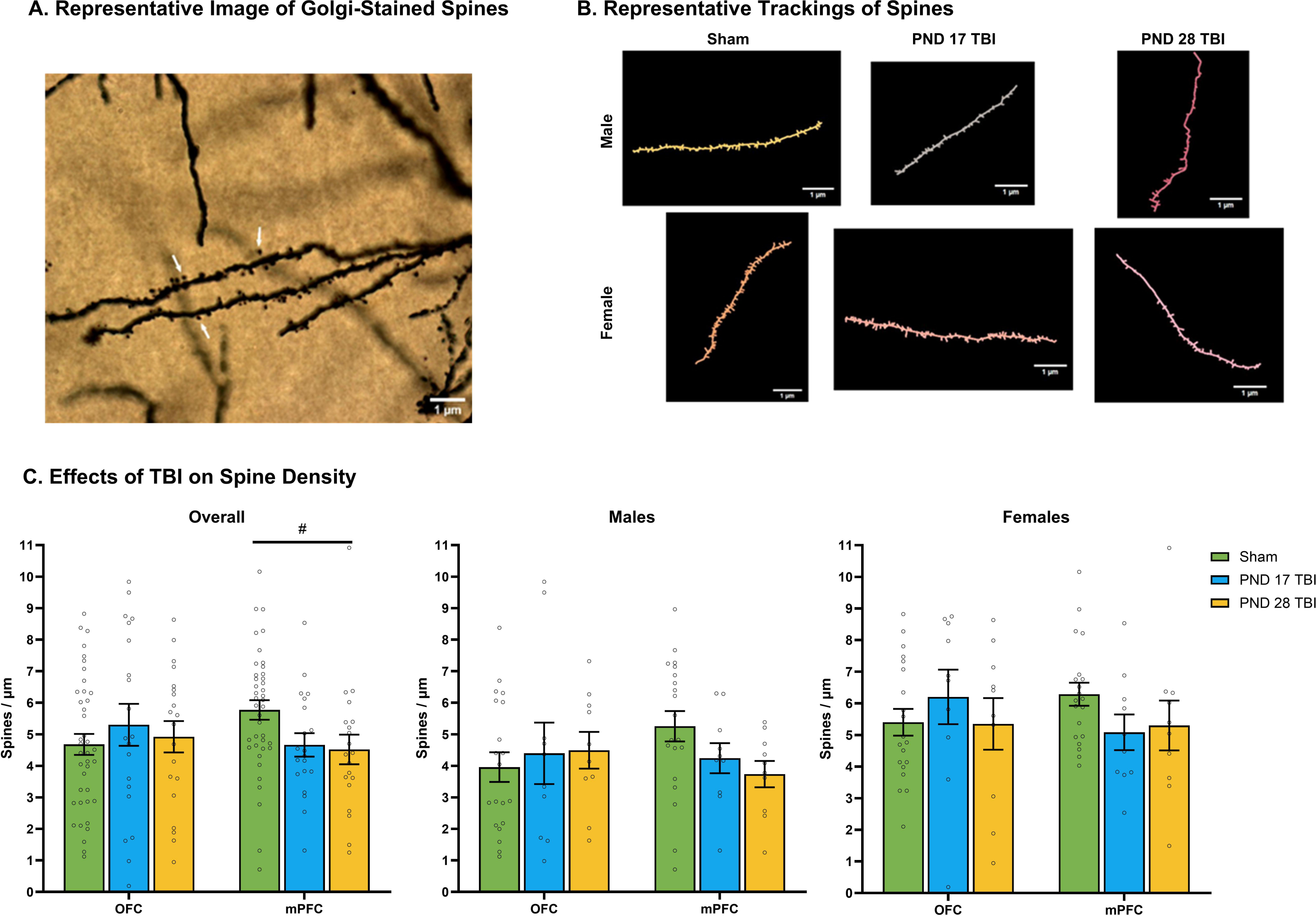
Effects of TBI on Spine Density in the Prefrontal Cortex. *Note.* Graphs show group means ± SEM. A-B) Scale bar = 1 µm. C) Neither TBI nor sex significantly impacted spine density in the mPFC or OFC, however, there was a large effect size in the mPFC, suggesting that TBI impacted spine density, but the sample size was too small. #, η^2^≥0.14

*Myelin ratio.* TBI also likely impacts the myelin levels in the OFC and mPFC as measured by calculating the myelin ratio (Figure 10). Our results indicate that a TBI on PND 17 decreased myelin while one on PND 28 increased myelin within the OFC. mPFC results display a similar trend where PND 17 TBI decreased myelin, and while PND 28 TBI myelin levels trended towards increased levels, it was not as drastically larger than sham animal as it was in the OFC. These trends do not support our hypothesis where we expected decreased myelin in the groups injured on PND 28. Regardless, this data suggests that a PND 17 frontal TBI imparts global deficit trends within the frontal lobe and neither injury affects distal brain regions.

**Figure 10.**
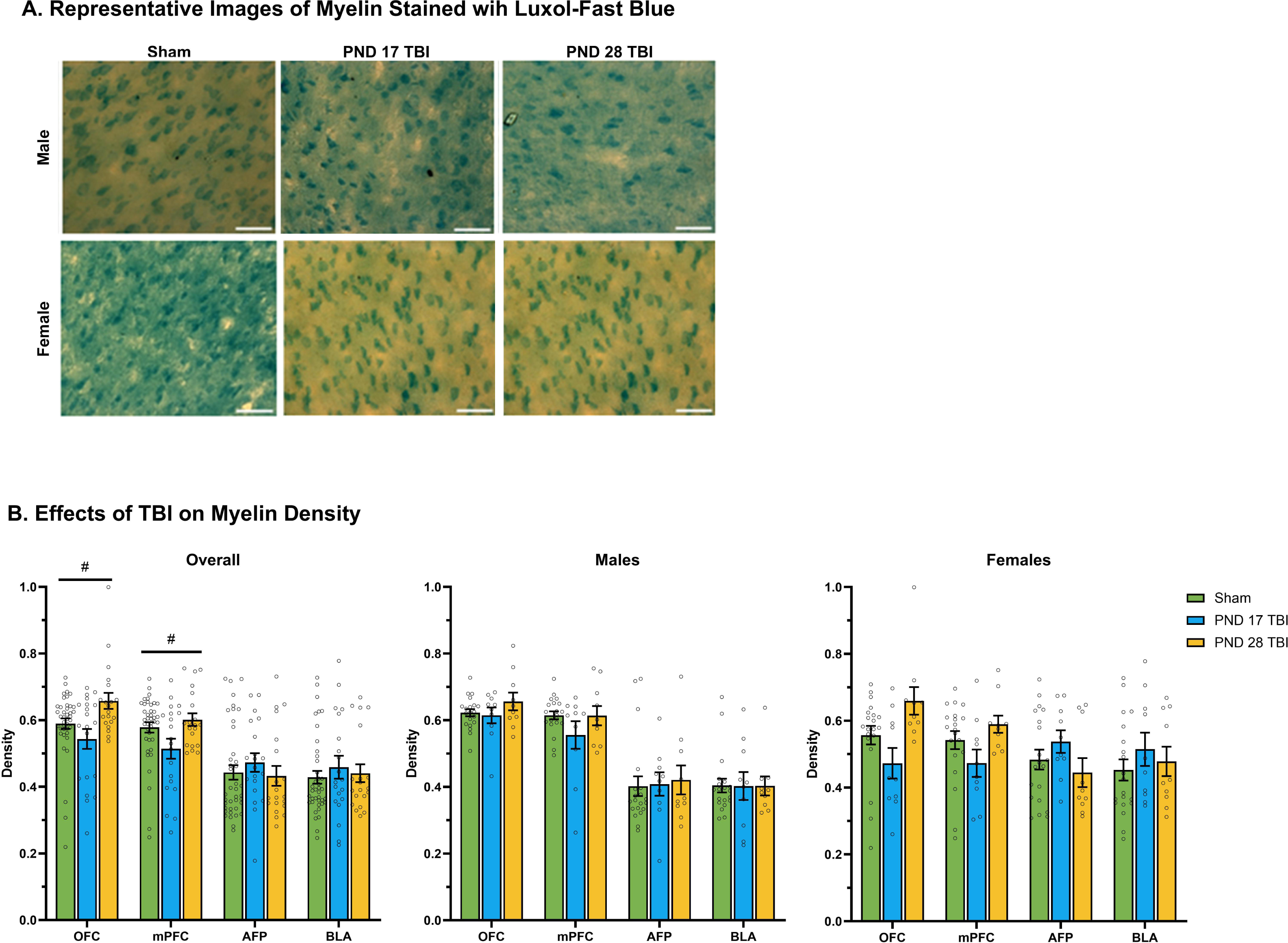
Effects of TBI on Myelin Density in the Prefrontal Cortex-Amygdala Pathway. *Note.* Graphs show group means ± SEM. A) Scale bar = 5 µm. B) Neither TBI nor sex significantly impacted myelin density in the mPFC or OFC, however, there was a large effect size in the mPFC and OFC, suggesting that TBI impacted myelin density in the PFC, but the sample size was too small. #, η^2^≥0.14

## Discussion

Overall, our data reveal two overarching patterns: (1) an age- and sex-specific effect of TBI on behavioral outcomes, and (2) a pronounced effect of PND 28 TBI on dominance and aggression. In the first pattern, females injured on PND 17 and males injured on PND 28 exhibited similar deficits across development, including impaired play behavior and heightened resident aggression. Whereas in the second pattern, PND 28 TBI, regardless of sex, increased dominance and intruder aggression. These patterns likely stem from three mechanisms: (1) age- and sex-specific neuroprotective mechanisms, (2) executive dysfunction, and (3) disrupted threat perception.

### Neuroprotection is Age- and Sex-Specific

Prior research has consistently shown greater neuroprotection following injury in females compared to males (7,55) that can be attributed to the neuroprotective effects of gonadal hormones, including estrogen and progesterone (57,85). Furthermore, females typically mature faster than males (7,86,87), with puberty occurring around PND 35 in female rats and PND 45 in male rats (88), which partially explains the discrepancies in our findings. Female animals injured on PND 28 exhibited fewer deficits compared to male animals injured on the same day, possibly due to the influence of female gonadal hormones that peak around puberty. Notably, post-menopausal women and ovariectomized animals do not demonstrate the same level of neuroprotection (57,58), suggesting that puberal hormones, irrespective of age confer neuroprotective effects. Importantly, gonadal hormones have an organization effect in the mPFC, via enhanced dendritic pruning (89) and promoting spine density in the mPFC (90,91). While we did not find significant morphological results, our effect sizes indicate age-specific deficits related to TBI and increased sample sizes are necessary to elucidate whether there are also sex-specific effects.

Regardless, behavioral results indicate that females injured on PND 28 and males injured on PND 17 display the most sparing of function. This pattern is likely influenced by sex-specific developmental timelines and gonadal hormones. For instance, females injured on PND 28 were nearing puberty, which may have temporally aligned with the neuroprotective effects of estrogen and progesterone that begins elevating on PND 30 (92,93). Whereas the sparing of function observed in a PND 17 TBI in males could be a result of the compensatory mechanisms of their earlier developmental stage (13,42). Indeed, more sparing of function and neuroprotective mechanisms have been observed following a PND 10 or older frontal lobe injury (9,25). Meanwhile, male PND 28 TBI animals, being older and lacking the protective influence of gonadal hormones available to female PND 28 TBI animals (92,93) or the developmental compensatory mechanisms seen in male PND 17 TBI animals (47,50,94), exhibit more deficits. Leaving female PND 17 TBI animals with the poorest outcomes, possibly due to sex-specific developmental pruning of mPFC neurons whereby females lose more neurons than males (89,95,96). Therefore, TBI-induced neurodegeneration may exaggerate normal developmental pruning in females injured on PND 17 and thus exacerbate functional deficits while reduced gonadal hormone neuroprotection in males injured on PND 28 contributes to their poor functional outcomes.

### Age- and Sex-Specific Executive Dysfunction

The mPFC undergoes a prolonged developmental period (3,17,19) that mirrors the gradual maturation of executive functions (18), including working memory, cognitive flexibility, and inhibitory control. All of these are essential for social interaction (97,98) as the PFC plays a pivotal role in the social brain network (99–101), influencing social motivation and behavioral adaptability in social contexts (102,103). Since social behavioral development parallels executive function maturation (98,104–106), this network is vulnerable to dysfunction following injury (107). Indeed, executive dysfunction is common following TBI (27,108), especially frontal TBI (27), most notably in impulse control (32–36,109).

Anatomically, executive functions undergo non-continuous developmental growth spurts beginning at birth and ending in young adulthood (22). Within the context of our research, a PND 17 TBI would occur during the period where there are dramatic enhancements in inhibitory control (110–112) and theory of mind (22), while a PND 28 TBI occurs during rapid improvements in working memory (113,114) and cognitive flexibility (115). The development of these behaviors is critical for the development of social behaviors, thus the social deficits observed in our study likely stem from impaired executive functions.

Notably, behavioral deficits in play behavior were observed across development but in sex- and age-specific patterns. The sex-specific differences observed could partially be due to pairing dynamics as female rats have been found to be more exclusionary than male rats towards rats with TBI (49). This potentially explains our observed protections from altered play dynamics in the male TBI animals prior to puberty. Meanwhile, the elevated play attacks observed in female TBI animals could be due to impulse control deficits, especially since play fighting requires inhibitory control and cooperation for accurate execution (116,117) and these deficits are common following mPFC injuries (32–36,109). This is especially likely in PND 17 TBI animals as they were injured during the developmental period with the greatest enhancements of inhibitory control (110–112), thus potentially disrupting development of this behavior. Meanwhile, PND 28 TBI tended to decrease play attack behaviors. Given that this injury occurred during the developmental period for working memory and cognitive flexibility (113–115), it is possible that deficits in these executive functions contributed to the observed decreases in play attacks. Working memory and cognitive flexibility work in tandem to aid in perception taking and subsequently behavioral adaptation (97,98,106). Furthermore, the mPFC regulates sociability, fear, and anxiety-like behavior (102,103) and deficits in theory of mind reduce play engagement (118,119) and impair social information processing (120,121). Thus, a PND 28 mPFC injury may disrupt development of working memory and theory of mind, thereby diminishing sociability and social approach.

However, we also observed changes play attack frequency across development whereby TBI animals that displayed exaggerated play attacks at one time point tended to demonstrate the opposite at a future time point, and vice versa. Social behavior is inherently dynamic and adjusts in response to environmental feedback (49,104,105,122). In fact, children with TBI often face increased peer rejection, victimization, fewer mutual friends, reduced popularity, greater shyness, and social withdrawal (122). Similarly, both male and female animals with TBI experience heightened social rejection, but it is particularly exaggerated in females (49). Our findings in female TBI animals generally align with this. Female TBI animals exhibited an initial increase in play attacks prior to puberty, followed by significant decreases thereafter. Although we did not directly measure social rejection, this pattern may explain the decline in play initiations observed in female TBI animals.

### Age-Specific Threat Perception Dysfunction

Due to the disinhibition of pre-existing aggressive tendencies (123,124), aggression is commonly increased following TBI, resulting in poor social functioning (125). The increased dominance and intruder aggression observed in the PND 28 TBI animals supports this. Along with the observed correlations between pre-puberty play attack frequency and adult resident offensive aggression, whereby female PND 17 TBI animals displayed a negative correlation and male PND 28 TBI animals displayed a positive correlation. Thus, TBI on PND 28 increased aggression and dominance regardless of sex, but especially in males, while a TBI on PND 17 disrupted developmental aggression trends in females.

While both female PND 17 and male PND 28 TBI animals displayed altered aggression, male PND 28 TBI animals seemed to display a stronger relationship between jTBI and aggression across development. However, it is debatable whether play attacks are the preamble to adult aggressive behavior. In fact, the opposite relationship may be true whereby higher play frequencies in childhood predict future subordinate behavior (126) as demonstrated in female PND 17 TBI animals in this study. Therefore, while female animals display higher aggression levels following PND 17 TBI, developmental patterns remain intact, aligning with previous research demonstrating lower aggression levels in females following TBI (56). Meanwhile, male PND 28 TBI animals display a disruption to this developmental pattern, which aligns with previous research showing higher aggression in males post-injury (56,127). Interestingly, this result indicates that elevated play behavior is an early warning sign for increased aggression and can serve as an indicatory for lifelong social deficits (104,105,122) further implicating the PND 28 time period as one of heightened risk for dominance and aggression following TBI.

Indeed, only TBI on PND 28 elevated intruder aggression and dominance. This age-specific effect potentially stems from alterations in the connectivity between the mPFC and amygdala, a key region implicated in regulating aggression (128,129). This circuit works via feedforward inhibition of excitatory neurons (130) to determine social ranks (131,132) and threat detection (102,103,128,129). Since the projections from the mPFC to the BLA mature in adolescence (133,134), a later time point than other mPFC pathways, a PND 28 TBI could disrupt the maturation of this circuit, thus causing deficits in social rank determination and threat detection and increasing aggressive tendencies (135).

However, it is also possible that the valence feedback encoding performed by the mPFC (103,136) is disrupted and appropriate processing of behavioral and emotional contexts is impaired. Indeed, social context influences the role of mPFC subregions with stimulations to the prelimbic-BLA circuit decreasing sociability under fear- and anxiety-inducing contexts (103,136) while stimulation of the infralimbic-BLA circuit increased sociability under stress-free conditions (103). Thus, in the context of our study, an mPFC injury on PND 28 may disrupt normal signaling to mPFC subregions that allows for appropriate threat signal perception and subsequently dysregulates aggression within the situational context and increasing aggression and dominance. This is especially prominent for intruder offensive aggression as it is not normal to display increase territorialness when intruding in another’s territory (68,137). Therefore, PND 28 TBI may disrupt the contextual valence processing between the mPFC and BLA, subsequently dysregulating and increasing aggressive behavior. While our histological data does not support this, effect sizes indicate that decreased spine density and increased myelin in the mPFC may be contributing to elevated aggression. More research is required to fully elucidate the neurological origins of TBI-induced aggression.

## Conclusions

Our research underscores the sex- and age-dependent impact of TBI on social behavior development. In males, childhood injuries (i.e., PND 28) tend to yield more pronounced deficits compared to females, whereas toddlerhood injuries (i.e., PND 17) in females result in the most significant deficits throughout development, including play initiation in puberty and aggression in adulthood. This discrepancy can be attributed to the differing developmental timelines between sexes, with females maturing faster than males. Thus, female animals injured closer to puberty benefit from neuroprotection conferred by gonadal hormones (57,85). However, male animals injured at the same age lack such neuroprotection due to a later onset of puberty (57,58), leading to social dysfunction. Conversely, male animals injured in toddlerhood, at a relatively immature developmental stage compared to equivalently aged female animals (7,86,87), demonstrate more sparing of function. In contrast, younger female animals are outside the scope of such neuroprotective mechanisms and experience social deficits. This research further highlights the impact of frontal TBI on dominance and aggression. Regardless of sex, damage to the mPFC increased dominance and aggression in adulthood, potentially due to the mPFC’s role in social rank (67,131,132) and impulse control (32–36,109). Overall, our research contributes to the large body of work regarding age of injury and sparing of function (8,9,11,13,14,25,26,47,50,66, for review see 138) but emphasizes the impact of sex on specific behavioral outcomes.

## Author Contributions

Sophia Shonka and Michael Hylin: conception or design of the work. Sophia Shonka: acquisition, analysis, and interpretation of data. Sophia Shonka and Michael Hylin: drafting or substantially revising the work.

