## Supplementary Material for "Younger is Better But Only for Males: Social Behavioral Development Following Juvenile Traumatic Brain Injury to the Prefrontal Cortex"

***Behavioral Model Training and Coding.***

To code the play behavior observations, social preference, social memory, and resident/intruder behaviors the DeepLabCut program (2.3.0), Behavioral Observation Research Interactive Software (BORIS, 7.13), and the Simulated Behavioral Analysis (SIMBA, 1.55) program were used.

*DeepLabCut.* First, body parts were tracked using DeepLabCut [1,2]. Videos of the three-chamber task (social preference and memory) were captured from above the rat and apparatus. Eight body parts were labeled for social preference and social memory, including left ear, right ear, nose, middle of back, left side, right side, tail base, and tail tip. Videos of the play behavior observations and resident/intruder task were captured from the side. Twelve body parts were labeled, including nose, right ear, left ear, right front paw, left front paw, back, right side, left side, right back paw, left back paw, tail base, and tail tip. For training, we extracted frames from 10% of the videos per behavior task, then we manually labeled 20 frames for each video. For the play behavior, social preference, and social memory tasks, frames were taken from a total of 24 videos for each task, eight of those videos came from each time point, and six from each group. For the resident/intruder task, frames were taken from a total of eight videos, two from each group.

Next, 95% of those labeled frames were used for training. We used a ResNet-50-based neural network [3,4] with default parameters for at least 200,000 training iterations. We then validated our model by assessing our pixel error rate and adjusting the model as necessary using the following steps. A pixel error rate of 5 or below was considered acceptable for the 480x640 resolution videos (social preference and memory videos, 0.00002% error) and an error of 20 or below was considered acceptable for the 720x1280 resolution videos (play behavior observations and resident/intruder task videos, 0.00002% error). If the pixel error rate was above that, then outlier frames were extracted, labels were corrected, and these frames were merged with our model. The model was then retrained and validated. If the pixel error rate still remained above the accepted threshold, then 20 more frames were extracted and labeled for each video, added to the training dataset, and a new training iteration was conducted. This process would have been repeated until the pixel error rate reached acceptable levels. While models can be overtrained by repeating these steps excessively, our models only needed at most one extra adjustment to reach satisfactory pixel error levels. Of the models that needed extra adjustments, only the play behavior observation and resident/intruder task models required an additional 20 frames per video be extracted and labeled. This was likely due to the higher frame rate in these videos, and thus, the model required a higher percentage of labeled frames to adequately train.

*BORIS.* Once post-estimation models were validated, behaviors were coded using BORIS software [5] by two independent researchers blinded to the experimental manipulations after achieving and 80% inter-rater reliability or better with each other according to Pearson's correlation coefficient. The same videos utilized for DeepLabCut training were utilized for behavior coding. The assessed behaviors for each task are outlined below.

For play behavior observations, animals were assessed for play attack and defense behaviors. In a typical play session, a pair of rats will compete for access to their partner's nape (a play initiation or attack), and if they reach the name of their partner, the rat will then nuzzle their partner [6,7]. Following an attack, the goal of the partner is to protect their nape and can respond with a variety of defensive tactics including evasions (swerving, leaping, or running away), complete rotations (rolling over into a prone position which leads to the rat being 'pinned'), partial rotations (rolling onto their side), and horizontal rotations (both rats stand on their hind legs and 'fight' with their fore paws). If a rat does not want to play, it can also just ignore the play initiation with a non-response (turning its head away from their partner and not engaging in the play session).

The three-chamber task consisted of two phases: social preference and social memory. In the social preference phase, the right and left compartments contained either a novel rat or a novel object. For the social memory phase, these compartments contained either a new novel rat or the now familiar rat from the social preference phase. Testing rats were placed in the center compartment and allowed to freely explore for 10 minutes. Interactions with the barriers (nose-touching, sniffing, climbing) that held the novel rat, novel object, and familiar rat was measured.

Finally, the resident/intruder task is commonly used to investigate aggression levels and allows for the evaluation of both offensive aggression and defensive aggression since subject animals can serve as both the resident (offensive aggression) and intruder (defensive aggression) across trials [8]. Offensive aggression is characterized as being the initiator of a range of threatening behavioral displays before attempting to attack non-vulnerable targets. This includes lateral threat (arched back and exposure of the flank), upright posture (on hind legs facing the opposing rat), clinch attack (a struggle or scuffle that is at close quarters, rats are entangled and intertwined), keep down (holding or pinning down opponent), and chase behaviors. Defensive aggression is performed in response to an attack from another animals and is distinct in its behavioral expression and inhibitory controls. This includes flight, defensive upright posture (opposing position to the upright posture), submission (displaying of stomach, opposing behavior to clinch attack), and freezing behaviors. Both these forms of aggression differ from violence, a pathological form of offensive behavior that has no functional value and includes attacks to vulnerable body parts, however none of our animals exhibited violent behavior so we were not able to code or train our model with this behavior. Other behavior that we coded includes social exploration (climbing over, sniffing non-ano-genital areas, and other interactions between animals), ano-genital sniffing, and social grooming (grooming of opponent).

*SimBA*. Pose estimation data from DeepLabCut and behavior coding data from BORIS was then input into the SimBA program [9] for behavior model training. Body part label outliers based on movement and location were corrected based on the length of the rat (distance between nose and tail base labels), and labels outside of two standard deviations from the median were corrected to the previous reliable location. Then 20% of video frames were withheld from training (roughly 5,940 frames out of 29,700 total frames per video). We used the eXtreme Gradient boost machine model with 2000 estimators, a minimum of 1 sample for a leaf node, square root for the number of features to consider when looking for the best split, and an entropy criterion to measure the quality of each split. Next, the model was validated using probability plots that were created for each behavior outlined above. These plots were examined to find the minimum probability threshold that would exclude false positives from each video. The average minimum threshold across all training videos was used (Table S2). Additionally, a minimum bout length was determined to smooth the probability plot and prevent false negatives. The minimum bout length was determined from the BORIS annotations for each video utilized in training. The average minimum bout length across all training videos was used (Table S2). The minimum threshold score and minimum bout length were then used to code for the presence of a behavior in each frame of the video. This was then used to determine inter-rater reliability between the researcher and the computer model. An inter-rater reliability score of at least 80% was required using Pearson's correlation coefficient before the model was finalized (Table S2). The resident/intruder task model additionally required the use of mutual exclusivity rules so that multiple behaviors were not encoded as occurring simultaneously (Table S3) but no other models required the use of such rules.

***Golgi-Cox Staining Procedures***

Staining procedures are based on previous research [10]. Animals were sacrificed on PND 63 via transcardial perfusion of 300 mL ice-cold phosphate buffered saline (PBS). Brains were then removed, blocked into three pieces (frontal lobe, parietal and occipital lobes, and cerebellum) and stored in the dark in 20 mL Golgi-Cox solution for 14 days. The Golgi-Cox solution was prepared from three stock solutions: solution A included 5% solution of potassium dichromate in distilled water; solution B included 5% solution of mercuric chloride in distilled water; solution C included a 5% solution of potassium chromate in distilled water. First, solutions A and B were mixed to 50% concentration of each. Next, solution C was diluted to 40% in distilled water. Finally, the AB solution mixture was combined with the solution C dilution and stored in a glass-stoppered bottle.

Next, the Golgi-Cox solution was replaced with 30% sucrose for at least 2 days in the dark before sectioning. Brains were sectioned into 200 μm coronal slices using a vibratome. Slices were then placed on a 2% gelatinized microscope slide and placed in a humidity-controlled glass staining tray for staining.

For staining, the slides were then rinsed in distilled water for 1 minute followed by ammonium hydroxide for 30 minutes in the dark. Slides were then rinsed in distilled water for 1 minute before incubation in Kodak Fix for Film for 30 minutes in the dark. Slides were rinsed in distilled water for 1 minute and dehydrated in 50% ethanol (EtOH) for 1 minute, 70% EtOH for 1 minute, 95% EtOH for 1 minute, and two successive rinses in 100% EtOH for 5 minutes. Finally, slides were placed in two successive rinses of xylene for 15 minutes then cover-slipped with Histomount.

***Luxol-Fast Blue Staining Procedures***

To measure myelin, Luxol-fast blue stain was utilized. Animals were sacrificed on PND 63 via transcardial perfusion of 300 mL ice-cold PBS followed by 300 mL of 4% paraformaldehyde. Brains were then removed and post-fixed in 4% paraformaldehyde at 4ºC for 48 hours before being changed into a fresh 30% sucrose solution for cryoprotection prior to sectioning. Frontal and parietal lobes were sectioned into 40 μm coronal slices using a cryostat. Slices used for staining were selected and soaked in PBS for 10 minutes followed by 70% EtOH for 24 hours to remove any lipids that remained from cryosectioning. Next, slices incubated in Luxol-fast blue solution (0.1% Luxol-fast blue, 0.5% glacial acetic acid, 99.4% 95%-EtOH) at 56ºC for four hours. Slices were then washed in distilled water ($dH_2O$) and PBS for three minutes each. Slices were differentiated in lithium carbonate solution (0.05% lithium carbonate, 99.05% distilled water) for 30 seconds followed by 70% EtOH for 30 seconds and rinsed in subsequent five-minute PBS washes. The slices were then mounted on 1% subbed slides and left to dry overnight. Following staining, slices were dehydrated in sequential washes of 70% EtOH (3 minutes), 95% EtOH (5 minutes), 100% EtOH (10 minutes), and xylene (10 minutes). The slides were then cover-slipped using Histomount.

### Supplementary Tables

**Table S1. Summary of Sham Group Results.**

*Note.* AFP, amygdalofugal pathway, BLA, basolateral amygdala, DF, degrees of freedom utilized in t-test analyses, mPFC, medial prefrontal cortex, N, total group size utilized in the Mann-Whitney U analyses, OFC, orbitofrontal cortex, SP, social preference, SM, social memory

| Variable | Statistical Test | DF/N | Test Statistic | p-value |
|---|---|---|---|---|
| Day 1 Foot Faults | Mann-Whitney U Test | 40 | -1.099 | 0.272 |
| Day 3 Foot Faults | Mann-Whitney U Test | 40 | 0.571 | 0.568 |
| Latency to Buried Food | Mann-Whitney U Test | 40 | 1.028 | 0.304 |
| Water-Almond Dishabituation Score | T-Test | 38 | 0.533 | 0.597 |
| Almond-Banana Dishabituation Score | Mann-Whitney U Test | 40 | 0.839 | 0.402 |
| Banana-Social 1 Dishabituation Score | Mann-Whitney U Test | 40 | 0.690 | 0.490 |
| Social 1-Social 2 Dishabituation Score | Mann-Whitney U Test | 40 | -1.028 | 0.304 |
| Habituation Score | Mann-Whitney U Test | 40 | -0.784 | 0.433 |
| # of Attacks - Pre-puberty | Mann-Whitney U Test | 40 | 0.041 | 0.967 |
| # of Attacks - Puberty | Mann-Whitney U Test | 40 | 1.481 | 0.139 |
| # of Attacks - Adulthood | Mann-Whitney U Test | 40 | -1.616 | 0.106 |
| # of Non-responses - Pre-puberty | Mann-Whitney U Test | 40 | -0.424 | 0.672 |
| # of Non-responses - Puberty | Mann-Whitney U Test | 40 | 0.067 | 0.947 |
| # of Non-responses - Adulthood | Mann-Whitney U Test | 40 | 0.919 | 0.358 |
| SP Score - Pre-puberty | Mann-Whitney U Test | 40 | 0.828 | 0.414 |
| SP Score - Puberty | Mann-Whitney U Test | 40 | -1.921 | 0.056 |
| SP Score - Adulthood | Mann-Whitney U Test | 40 | -1.163 | 0.253 |
| SM Score - Pre-puberty | T-Test | 38 | 1.177 | 0.247 |
| SM Score - Puberty | T-Test | 38 | -0.577 | 0.284 |
| SM Score - Adulthood | T-Test | 38 | -1.050 | 0.3 |
| Number of Winning Trials in the Tube Task | Mann-Whitney U Test | 40 | -0.373 | 0.709 |
| Resident Offense Score | Mann-Whitney U Test | 40 | 0.082 | 0.935 |
| Intruder Offense Score | Mann-Whitney U Test | 40 | -1.145 | 0.252 |

| Variable | Statistical Test | DF/N | Test Statistic | p-value |
|---|---|---|---|---|
| Resident Violence Score | Mann-Whitney U Test | 40 | 0.344 | 0.731 |
| Intruder Violence Score | Mann-Whitney U Test | 40 | 0.296 | 0.767 |
| Critical value for OFC Basal Dendrites | T-Test | 38 | -1.120 | 0.135 |
| Dendrite Length for OFC Basal Dendrites | T-Test | 38 | -1.103 | 0.138 |
| Critical value for OFC Apical Dendrites | T-Test | 38 | 0.169 | 0.433 |
| Dendrite Length for OFC Apical Dendrites | Mann-Whitney U Test | 40 | -0.081 | 0.935 |
| Critical value for mPFC Basal Dendrites | T-Test | 38 | 1.051 | 0.15 |
| Dendrite Length for mPFC Basal Dendrites | T-Test | 38 | -0.504 | 0.309 |
| Critical value for mPFC Apical Dendrites | T-Test | 38 | -1.424 | 0.081 |
| Dendrite Length for mPFC Apical Dendrites | T-Test | -1.061 | 38 | 0.148 |
| Spine Density - mPFC | Mann-Whitney U Test | 40 | -0.135 | 0.892 |
| Spine Density - OFC | Mann-Whitney U Test | 40 | 0.054 | 0.957 |
| Myelin Ratio - mPFC | Mann-Whitney U Test | 40 | -1.001 | 0.317 |
| Myelin Ratio - OFC | Mann-Whitney U Test | 40 | 0.162 | 0.871 |
| Myelin Ratio - AFP | Mann-Whitney U Test | 40 | 0.108 | 0.914 |
| Myelin Ratio - BLA | Mann-Whitney U Test | 40 | 1.407 | 0.160 |

**Table S2. Reliability and Validation of Behavior Models**

*Note.* Inter-rater reliability is between the AI model and the researcher and was averaged across the videos that were utilized for training. *, behavior classification was validated but not utilized in the current study.

| Mod el | Behavior | Body Labels Error Rate | Minimum Probability Threshold | Minimum Bout Length | Inter-rater Reliability |
|---|---|---|---|---|---|
| Social Preference | | 2.68 pixels | | | |
| | Novel Object Interactions | | 0.4395 | 1 second | 0.984 |
| | Novel Rat Interactions | | 0.6492 | 1 second | 0.98 |
| Social Memory | | 2.53 pixels | | | |
| | Familiar Rat Interactions | | 0.6352 | 1 second | 0.979 |
| | Novel Rat Interactions | | 0.5374 | 1 second | 0.982 |
| Play Behavior Observations | | 11.35 pixels | | | |
| | Play Attack | | 0.465 | 115 ms | 0.938 |
| | Nonresponse | | 0.4 | 185 ms | 0.929 |
| Resident/Intruder Task | | 19.23 pixels | | | |
| | Lateral Threats | | 0.3428 | 34 ms | 0.954 |
| | Upright Posturing | | 0.3682 | 325 ms | 0.921 |
| | Clinch Attack | | 0.4476 | 25 ms | 0.938 |
| | Keep Down | | 0.4109 | 232 ms | 0.915 |
| | Chase | | 0.2864 | 560 ms | 0.921 |
| | *Social Exploration | | 0.4158 | 131 ms | 0.935 |
| | *Ano-Genital Sniffing | | 0.5253 | 102 ms | 0.936 |
| | *Social Grooming | | 0.4180 | 914 ms | 0.881 |
| | *Flight | | 0.1994 | 334 ms | 0.923 |
| | *Defensive Upright Posture | | 0.4243 | 241 ms | 0.923 |
| | *Submission | | 0.4781 | 288 ms | 0.942 |
| | *Freeze | | 0.6138 | 275 ms | 0.926 |

**Table S3. Mutual Exclusivity Rules for Resident/Intruder Task**

| Behavior 1 | vs. | Behavior 2 | Winner |
|---|---|---|---|
| Social Exploration | | Lateral Threat | Social Exploration |
| Social Exploration | | Ano-Genital Sniffing | Ano-Genital Sniffing |
| Social Exploration | | Upright Posture | Upright Posture |
| Social Exploration | | Clinch Attack | Clinch Attack |
| Social Exploration | | Chase | Social Exploration |
| Social Exploration | | Social grooming | Social Grooming |
| Social Exploration | | Defensive Upright Posture | Social Exploration |
| Ano-Genital Sniffing | | Social Grooming | Ano-Genital Sniffing |
| Lateral Threat | | Social Grooming | Social Grooming |
| Lateral Threat | | Freeze | Freeze |
| Upright Posture | | Clinch Attack | Upright Posture |
| Upright Posture | | Defensive Upright Posture | Highest Threshold |
| Keep Down | | Chase | Chase |
| Keep Down | | Social Grooming | Social Grooming |
| Keep Down | | Defensive Upright Posture | Keep Down |
| Defensive Upright Posture | | Submission | Defensive Upright Posture |
| Defensive Upright Posture | | Freeze | Freeze |

### Supplemental Figures

**Figure S1. Effects of TBI on Performance in the Three Chamber Task.**

*Note.* Graphs show group means ± SEM. A) TBI did not significantly impact social preference. B) TBI did not significantly impact social memory.

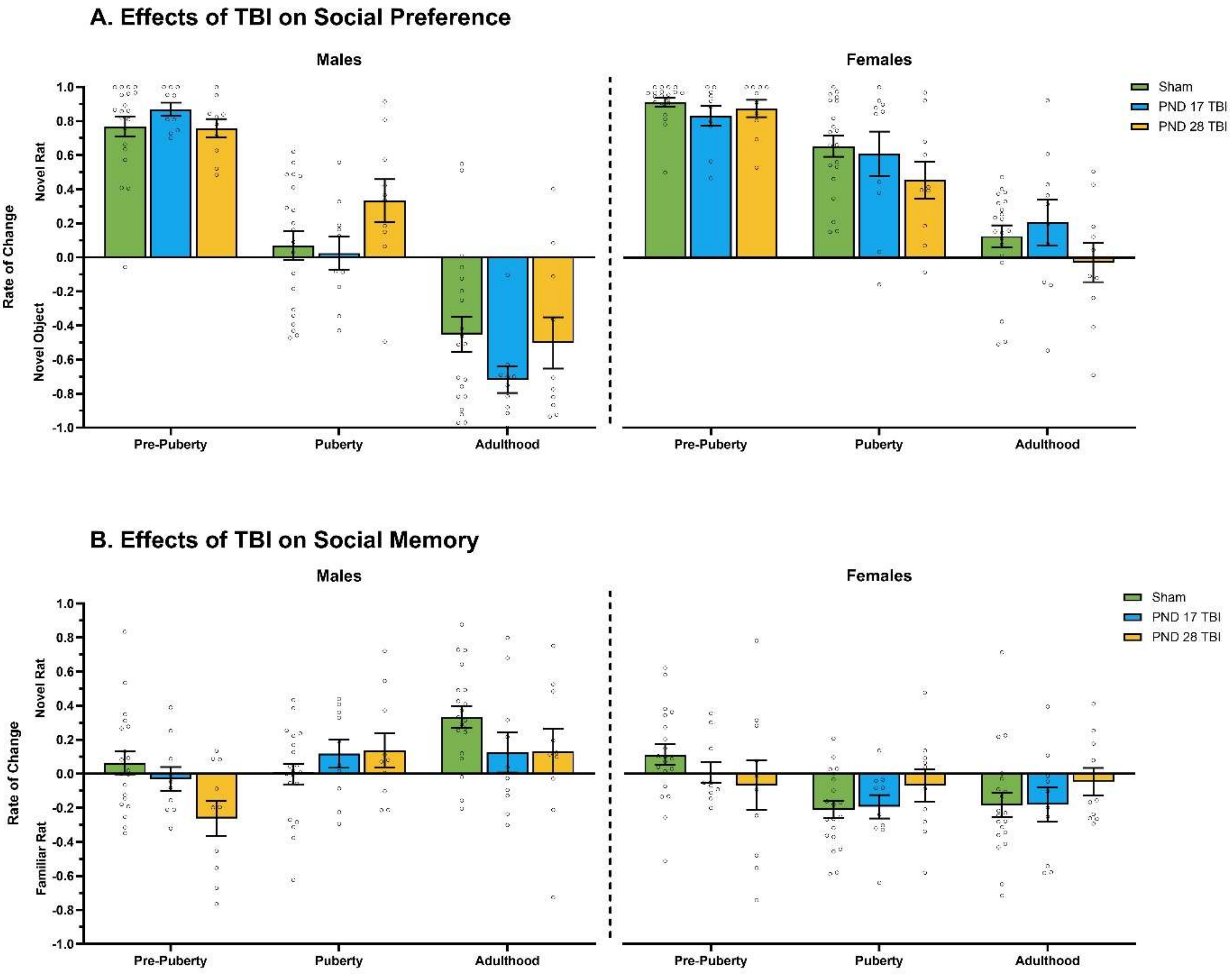

**Figure S2. Impact of Childhood Play Behavior on Adult Social Dominance.**

*Note.* Graphs show individual data points with linear regression lines. There was no correlation between childhood play behavior and adult dominance behaviors, and neither TBI nor sex moderated that relationship.

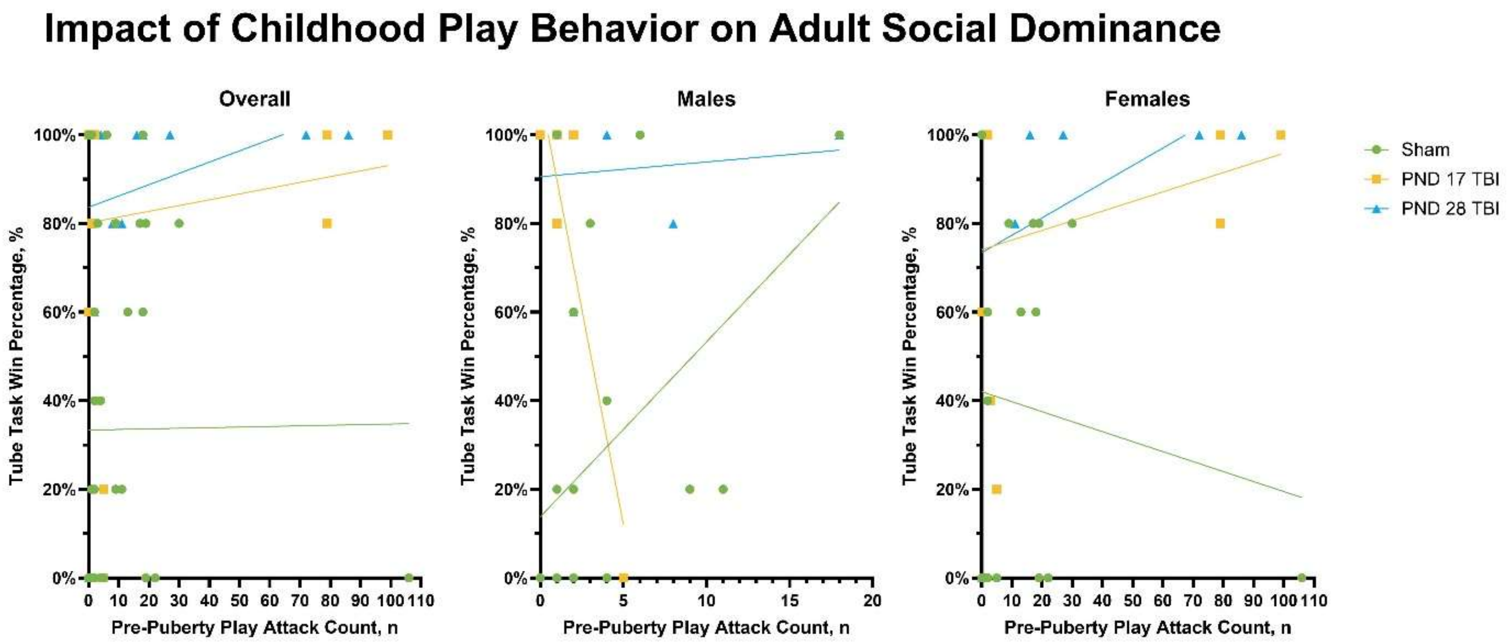

**Figure S3. Impact of Childhood Play Behavior on Adult Violence in the Resident/Intruder Task.**

*Note.* Graphs show individual data points with linear regression lines. A-B) There was no significant effect of childhood play behavior on adult violence in either the resident or intruder phase and neither TBI nor sex moderated this relationship.

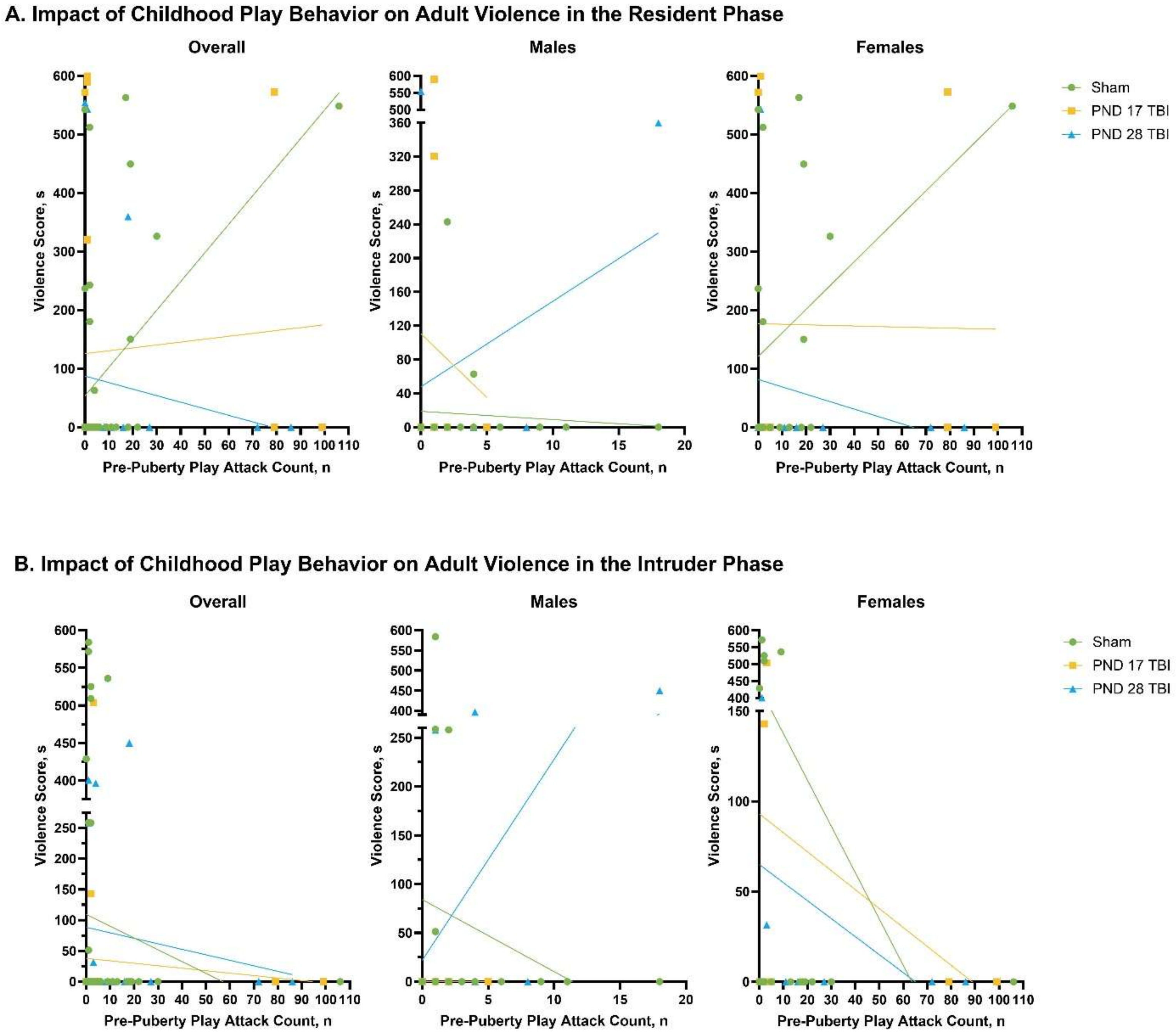
